# A mechanism for hippocampal memory recall based on excitatory-inhibitory fluctuations in neocortex

**DOI:** 10.1101/2020.11.27.401299

**Authors:** Renée S. Koolschijn, Anna Shpektor, I. Betina Ip, William T. Clarke, David Dupret, Uzay E. Emir, Helen C. Barron

## Abstract

The brain has a remarkable capacity to acquire and store memories that can later be selectively recalled. These processes are supported by the hippocampus which is thought to index memory recall by reinstating information stored across distributed neocortical circuits. However, the mechanism that supports this interaction remains unclear. Here, in humans, we show that recall of a visual cue from a paired associate is accompanied by a transient increase in the ratio between glutamate and GABA in visual cortex. Moreover, these excitatory-inhibitory fluctuations are predicted by activity in the hippocampus. These data suggest the hippocampus gates memory recall by indexing information stored across neocortical circuits using a disinhibitory mechanism.

## INTRODUCTION

Memories are thought to be stored across sparse and distributed neuronal ensembles in the brain^1,2^. To facilitate memory recall, activity across neuronal ensembles is selectively reinstated to recover enduring representations of the past. This reinstatement is thought to be mediated by the hippocampus, a brain region important for learning and memory^3^. Anatomically, the hippocampus sits at the apex of a cortical sensory processing hierarchy^4^ where inputs received by sensory cortices reach the hippocampus via the entorhinal cortex and other relay regions, which in turn make widespread cortico-cortical connections that project the hippocampal output back to neocortex^5,6^. This reciprocal anatomical connectivity equips the hippocampus with the necessary architecture to coordinate activity with neocortex, thus providing a ‘memory index’, or summary sketch, for information stored across distributed cortical circuits^7–9^. Consistent with this view, during memory recall hippocampal reinstatement predicts subsequent neocortical reinstatement^10^.

However, the mechanism that allows the hippocampus to coordinate reinstatement across distributed neocortical circuits remains unclear. In animal models, neural circuit manipulations suggest higher-order brain regions may modulate release of sensory information in neocortex via disinhibitory circuit mechanisms^11,12^. For example, during attentional modulation, projections from the cingulate region of mouse frontal cortex modulate GABAergic circuits in visual cortex to enhance visual discrimination^13^. Building upon this idea, one possibility is that the hippocampus mediates memory recall using a similar mechanism, by transiently modulating the relationship between neocortical excitation and inhibition.

At the cellular level, tight coupling between neocortical excitation and inhibition (EI) can be observed during both sensory stimulation and spontaneous neural activity^14–16^. This phenomenon has led to the concept of EI balance, where, following changes in excitability, synaptic strength, current or overall network activity returns to a stable set point via negative feedback^17^. Therefore, while microcircuits are capable of large changes in activity due to synaptic delays or differences in signal propagation speed, transient excitatory responses are rapidly quenched by inhibition^16,18^. The dynamic interplay between excitation and inhibition may therefore shape computations performed by cortical circuits, including in response to inputs that derive from brain regions such as the hippocampus.

While physiological measures of EI balance vary in both definition and granularity, non-invasive methods available for imaging the human brain are acquired at a more coarse spatiotemporal scale. Magnetic Resonance Spectroscopy (MRS) provides a unique tool to quantify the concentration of different neural metabolites^19,20^, including glutamate and GABA, the principle excitatory and inhibitory neurotransmitters in the brain. While MRS cannot dissociate between neurotransmitter and metabolic pools of glutamate and GABA^21,22^, meaningful interpretation of MRS-derived measures derives from a major body of work showing an approximately 1:1 relationship between the rate of glutamine-glutamate cycling, which is necessary for glutamate and GABA synthesis, and neuronal oxidative glucose consumption, which indirectly supports neurotransmitter release among other processes^23–25^. Moreover, despite providing an indirect measure, MRS-derived glutamate and GABA reported during learning and memory paradigms in humans show remarkable correspondence with findings reported at the physiological level in animals. For example, in animals a reduction in GABAergic tone is necessary for induction of neocortical plasticity via long-term potentiation (LTP)^26,27^, while in humans motor learning and plasticity in visual cortex are accompanied by a reduction in MRS-derived GABA^28,29^. Investigations in both animal models and humans further show that after new learning EI balance prevails to ensure memories are stored in a stable and dormant state^30–32^. This leads to the following prediction: memory recall involves a transient break in EI balance, opening a window to release memories from the blanket of inhibition before re-establishing network stability. Moreover, MRS-derived measures of glutamate and GABA may provide a suitable index for this process.

To test this prediction, here we implement a new sequence that combines functional Magnetic Resonance Imaging (fMRI) with functional MRS (fMRS). Together with an event-related design, we use the Blood-Oxygen-Level-Dependent (BOLD) signal to probe hippocampal-dependent associative memory recall of a visual cue, while simultaneously measuring dynamic changes in MRS-derived glutamate and GABA in visual cortex. During memory recall, we report a transient increase in the ratio between MRS-derived glutamate and GABA in neocortex which is selectively predicted by the BOLD signal in the hippocampus. These findings suggest the hippocampus indexes recall by transiently modulating neocortical EI balance to release memories stored across distributed neural circuits.

## RESULTS

### Task design and behaviour

To investigate the neuronal mechanisms that support memory recall we designed a three-stage inference task (Fig. 1a) that has previously been shown to involve associative memory recall in humans and mice^33^. Moreover, we chose to implement an inference task because, unlike some forms of first-order associative recall, previous lesion and optogenetic studies in rodents demonstrate that second-order associative recall required for inference is a hippocampal dependent process^33–35^. Thus, the inference task provided an opportunity to investigate how the hippocampus mediates neocortical excitation and inhibition during memory recall.

**Figure 1.**
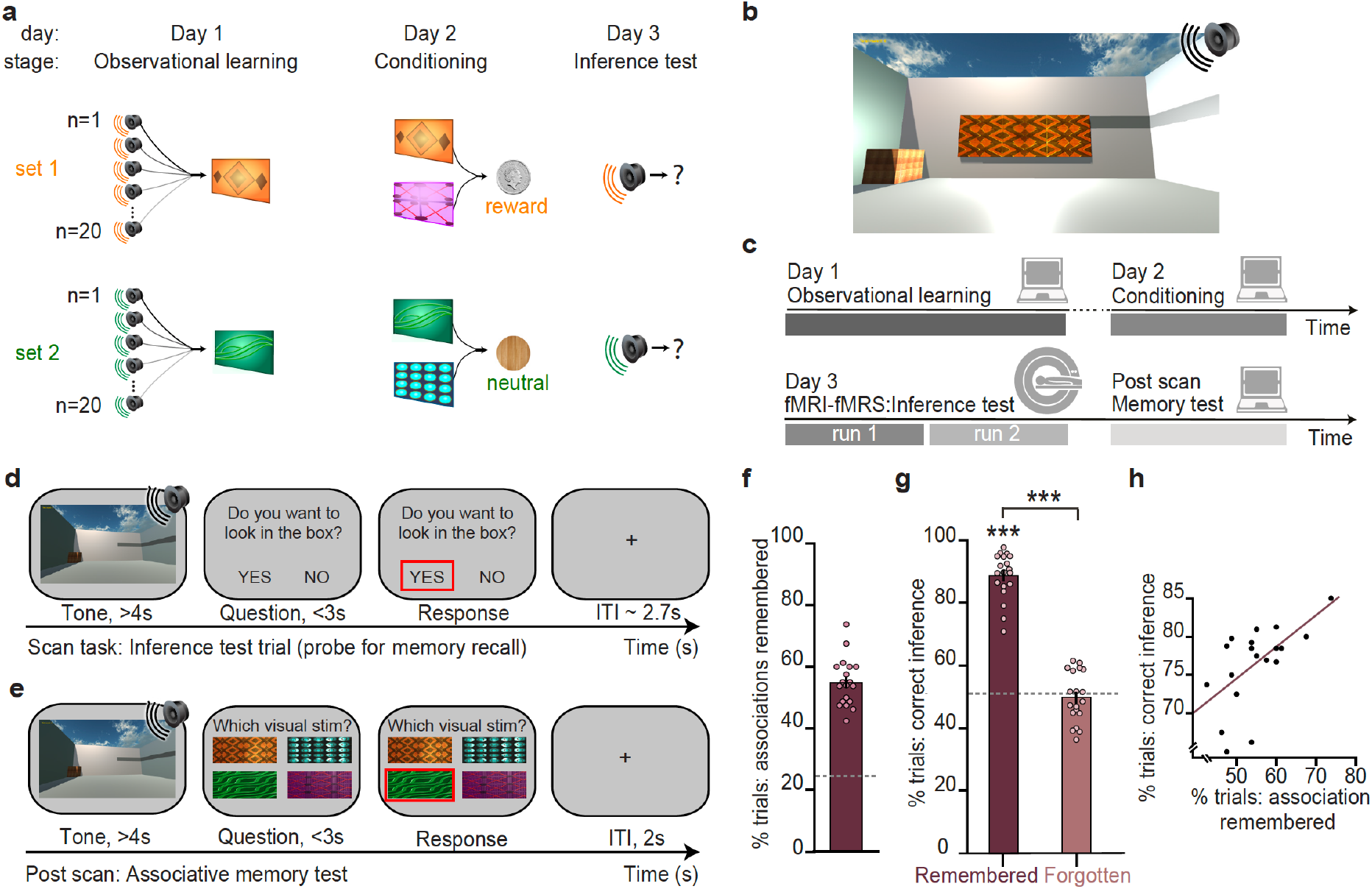
Inference task design and behavioural performance. **a** Three-stage inference task designed to investigate hippocampal-dependent associative memory recall. First, participants learned to associate auditory cues with visual cues (‘observational learning’ stage, day 1), where four different visual cues were each associated with 20 auditory cues. Second, participants learned to associate visual cues with an outcome (‘conditioning’ stage, day 2), where two visual cues predicted a rewarding outcome (set 1, monetary coin) while the other two predicted a neutral outcome (set 2, woodchip). Third, the auditory cues were played in isolation and we assessed participants’ ability to infer the relevant outcome (‘inference test’, day 3). **b** The three-stage inference task shown in *a* was performed within a virtual-reality environment. **c** Schematic: training and testing protocol. The inference test was performed inside the 7T MRI scanner. After completion of the three-stage inference task participants were given a surprise memory test (day 3). **d** Example inference test trial performed inside the scanner. For each auditory cue, participants were required to infer whether they would like to look in the wooden box, where the outcome cues were delivered during conditioning. **e** Example trial from the surprise post-scan associative memory test. **f** During the post-scan associative memory test participants remembered 55% of the auditory-visual associations (54.8 ±1.78%; mean ±SEM), significantly above chance as indicated by the dotted line (t_18_=16.80 p<0.001). **g** Behaviour during the inference test (Fig. 1d) was assessed as correct if participants pressed ‘yes’ for auditory cues in set 1, or ‘no’ for auditory cues in set 2. Participants successfully inferred on trials for which the auditory-visual association was later remembered (‘later remembered’: t_18_=22.91, p<0.001; ‘later forgotten’: t_18_=0.09, p=0.925; ‘later remembered’ – ‘later forgotten’: t_18_=16.21, p<0.001; dotted line indicates chance). **h** Across participants, behavioural performance on the inference test was predicted by behavioural performance on the post-scan associative memory test (r_17_=0.57, p=0.010). Notably, there was no significant effect of sex on behavioural performance (Supplementary Table 1).

In the first stage of the task participants learned up to 80 auditory-visual associations (‘observational learning’, day 1; Fig. 1a, Supplementary Fig. 1). In the second stage, which occurred approximately 24 hours later, each visual cue was paired with either a rewarding (set 1) or neutral outcome (set 2) (‘conditioning’, day 2; Fig. 1a, Supplementary Fig. 1). Rewarding outcomes were silver coins that were later exchangeable for a monetary sum, while neutral outcomes were non-exchangeable woodchips. Importantly, auditory cues were never paired with an outcome, providing an opportunity to assess evidence for an inferred relationship between these indirectly related stimuli. Accordingly, in the third stage of the task we presented auditory cues in isolation, without visual cues or outcomes, and we measured evidence for inference from the auditory cues to the appropriate outcome (‘inference test’, day 3; Fig. 1a). All stages of the task, including the day 3 inference test were performed in virtual reality (VR) (Fig. 1b), an immersive and highly controlled 3D environment that has the potential to benefit from cross-species comparisons in the future^33^.

Participants performed the day 3 inference test during an MRI scan (Fig. 1c–d). In response to the auditory cues in the inference test, participants successfully inferred the correct outcome if they could later recall the relevant auditory-visual association during a surprise post-scan associative memory test performed after the inference task was completed (Fig. 1e–g). Indeed, performance on the associative memory test, that assessed memory for auditory-visual associations learned on day 1, predicted performance on the inference test performed on day 3 (Fig. 1h). Consistent with previous neuroimaging data in humans and cellular recordings in mice^33^, these behavioural findings suggest inferential choice during the inference test involves associative recall of the intermediary visual cues. In this manner, the inference task provides a suitable paradigm to investigate the neural mechanisms that underlie hippocampal-dependent associative memory recall.

### BOLD signal in the hippocampus and visual cortex is modulated during memory recall

To investigate the relationship between the hippocampus and neocortex during associative memory recall we implemented a novel imaging sequence^36^, which enabled interleaved acquisition of near-whole brain fMRI together with fMRS in V1 (Fig. 2a). This imaging sequence thus provided a means to simultaneously measure both hemodynamic and neurochemical changes during the inference task, in an event-related manner.

**Figure 2.**
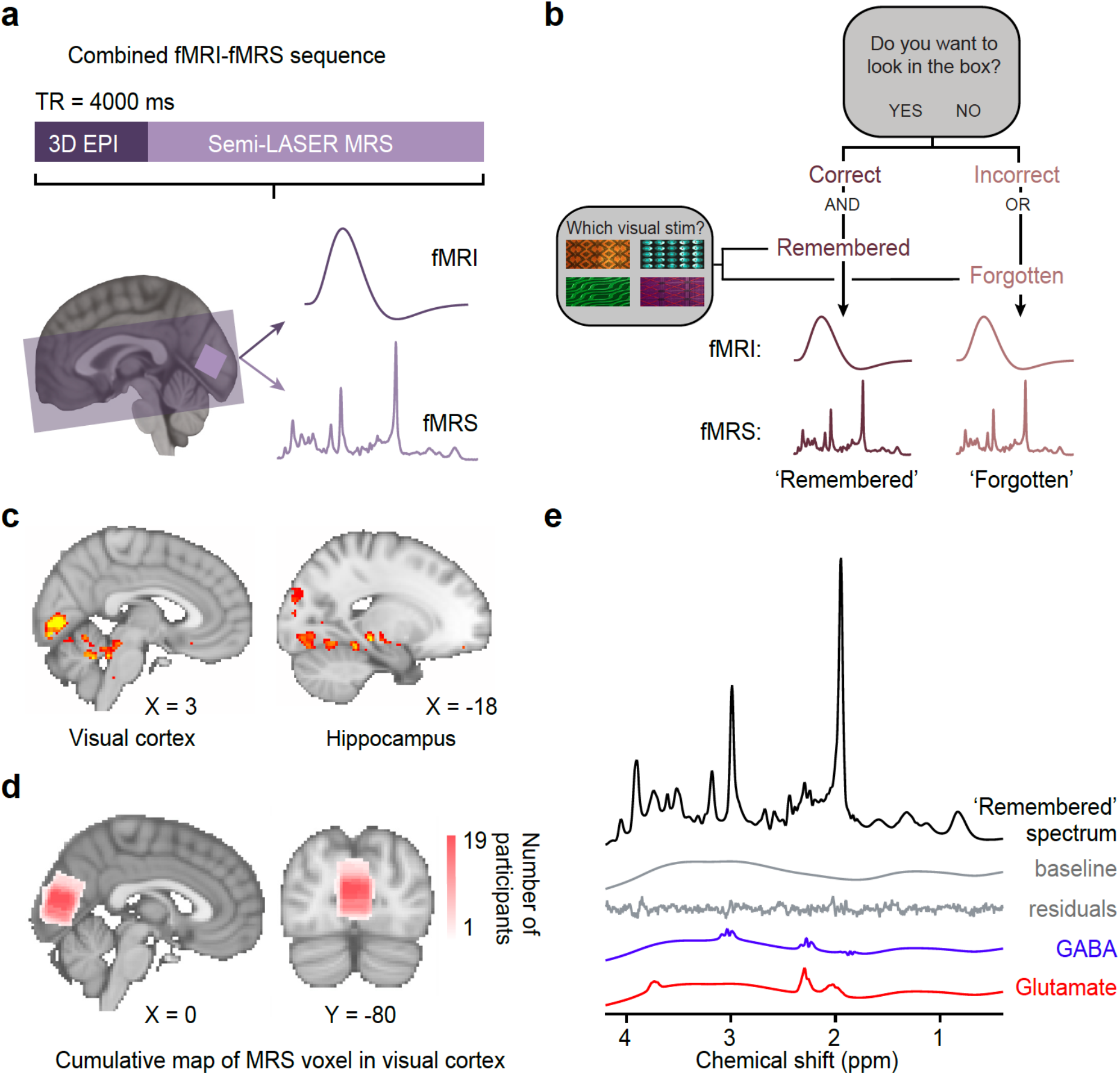
Using fMRI-fMRS data to assess changes in BOLD signal and glu/GABA ratio during the inference test. **a**7T MRI sequence. 3D BOLD echo planar imaging (3D-EPI) and semi-LASER MR-spectroscopy were acquired in the same TR. The MRS voxel was positioned in V1 (light-purple) and the EPI slice coverage included occipital and temporal lobes (dark-purple). **b** Schematic showing how trials during the inference test were categorized into ‘remembered’ and ‘forgotten’. Trials were categorised as ‘remembered’ if participants correctly inferred the appropriate outcome during the inference test *and* subsequently recalled the auditory-visual association in the post-scan memory test. Trials were categorised as ‘forgotten’ if participants incorrectly inferred the appropriate outcome during the inference test *or* subsequently forgot the auditory-visual association in the post-scan memory test. **c** During the question period in the inference test (Fig. 1c–d), BOLD signal in the visual cortex and the hippocampus was significantly higher for ‘remembered’ versus ‘forgotten’ auditory cues (‘remembered’ – ‘forgotten’, visual cortex: t_17_=6.93, p<0.001; left hippocampus: t_17_=4.36, p=0.017; whole-volume FWE-corrected; together with regions listed in Supplementary Table 3). **d** Anatomical location of 2×2×2 cm^3^ MRS voxel positioned in V1. Cumulative map across participants. **e** Representative MRS spectrum from ‘remembered’ trials in the inference test, for an example subject. Top to bottom: average spectra, baseline, residuals, estimated GABA, estimated glutamate.

Using fMRI data from the interleaved sequence, we first identified brain regions modulated by recall of a visual cue in response to the associated auditory cue presented during the inference test (Fig. 1a). To obtain the most accurate estimate for associative memory recall, we categorized trials post-hoc, using participants’ behavioural performance from both the inference test and subsequent post-scan associative memory test (Fig. 1c–e), which were highly correlated across participants (Fig. 1h). Trials where participants made both the correct inference and subsequently remembered the auditory-visual associations were classified as ‘remembered’. Trials where participants made either the incorrect inference or subsequently forgot the auditory-visual associations were classified as ‘forgotten’ (Fig. 2b, Supplementary Table 2, *Methods*). Neural signatures acquired during the ‘forgotten’ trials thus provided a control condition for those acquired during the ‘remembered’ trials. Consistent with previous research investigating associative recall of visual cues^37,38^, we observed a significant increase in BOLD signal in both the hippocampus and visual cortex on ‘remembered’ versus ‘forgotten’ trials (Fig. 2c; Supplementary Fig. 2).

### Dynamic increase in the ratio between glutamate and GABA in visual cortex during recall

We then asked whether associative memory recall of a visual cue is also accompanied by changes in the ratio between glutamate and GABA (‘glu/GABA ratio’) in the visual cortex. Using the interleaved fMRS data acquired in primary visual cortex (V1) (Fig. 2a,d), we quantified the concentration of glutamate and GABA normalised to total Creatine (tCr) in an event-related manner (Fig. 2b,e). We then used MRS-derived measures of glutamate and GABA to estimate changes in glu/GABA ratio^39^ (see *Methods*), where changes are evaluated through assessment of the ratio of ‘remembered’ trials relative to ‘forgotten’ (as defined above). In this manner, the ‘forgotten’ trials again provide a condition and stimulus-matched control for data acquired during the ‘remembered’ trials.

To detect dynamic changes in glu/GABA ratio it was not appropriate to implement default assumptions typically used to detect static estimates (see *Methods*). Namely, these default assumptions assume the dynamic range of GABA is fixed by normalising GABA relative to other more abundant metabolites. Here, to optimise our sensitivity to changes in glu/GABA across conditions we removed these default constraints. Notably, while this approach leads to higher GABA estimates, the uncertainty in the metabolite estimates were reduced (Supplementary Fig. 3). Moreover, our analysis controlled for any effect of metabolite scaling by comparing the difference between two conditions (‘remembered’ versus ‘forgotten’).

During recall, we observed an increase in glu/GABA ratio in V1 when comparing ‘remembered’ versus ‘forgotten’ cues (Fig. 3a–b). Standard quality metrics indicated that our data quality was comparable with those reported in previous studies^40–43^ (Supplementary Fig. 4, Supplementary Table 4). To control for any biases introduced by differences in the number of ‘remembered’ versus ‘forgotten’ trials (Supplementary Table 5), we compared the group mean metabolite change against a null distribution generated by permuting the identity labels assigned to each trial. This analysis revealed a significant decrease in GABA and a significant increase in glu/GABA ratio during memory recall (Fig. 3d–f). This change in glu/GABA ratio was still observed when using performance on the inference task alone to categorise trials into ‘remembered’ and ‘forgotten’ (Supplementary Fig. 5). Furthermore, the increase in glu/GABA ratio was not observed during periods immediately before or after recall (Fig. 3a–b; Supplementary Fig. 6). These findings cannot be explained by differences in data quality measures between the ‘remembered’ and ‘forgotten’ conditions (Supplementary Fig. 7). Moreover, no effect was observed in NAA, which has overlapping peaks with GABA but is found at higher concentration (Supplementary Fig. 8). Thus, we propose this transient increase in neocortical glu/GABA ratio reflects a mechanism for associative memory recall.

**Figure 3.**
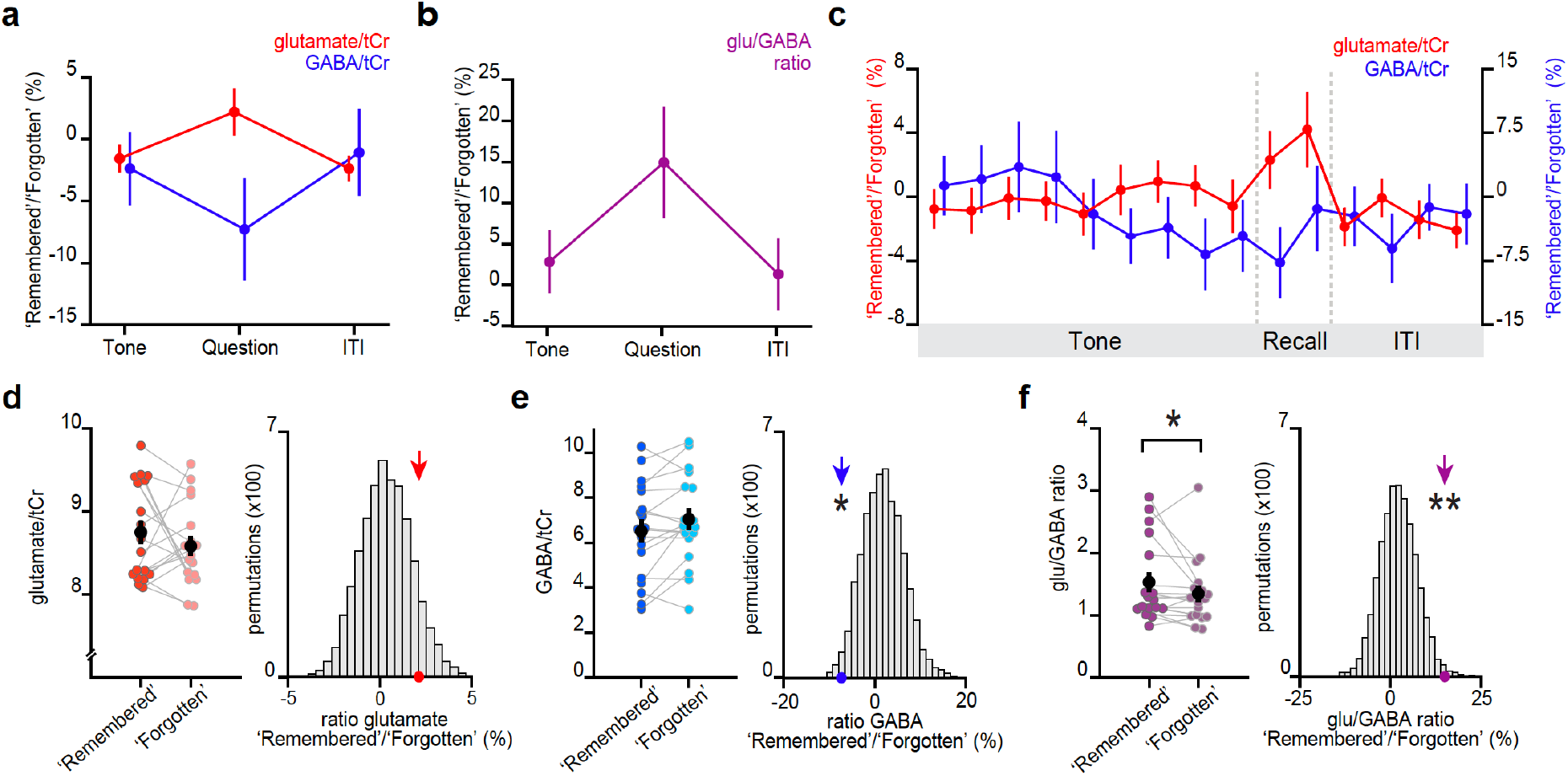
Memory recall and inference involves a transient break in glu/GABA ratio. **a-b** During the question period of the inference test trials (up to 3 s), glu/GABA ratio significantly increased during ‘remembered’ versus ‘forgotten’ trials (‘remembered’:‘forgotten’, glu/GABA ratio: t_17_=2.19, p=0.042). This break in glu/GABA ratio was not observed during the ‘tone’ (~7 s) or ‘ITI’ (~2.7 s) periods (‘Tone’, glu/GABA ratio: t_18_=0.74, p=0.468; ‘ITI’, glu/GABA ratio: t_18_=0.30, p=0.766). Note that glutamate:tCR and GABA:tCr concentrations have been multiplied by 8 as per LCModel’s default settings. **c** Moving average showing glutamate:tCr and GABA:tCr for the ratio of ‘remembered’ to ‘forgotten’ trials during the inference test. Each point represents a 2.5s time bin (mean ± SEM). **d-f** Left: The metabolite values and glu/GABA ratio during the question period for ‘remembered’ and ‘forgotten’ trials (mean ± SEM). Right: Comparing the mean ratio of ‘remembered’ to ‘forgotten’ (coloured arrows) against null distributions generated by permuting the trial labels to control for any potential biases in the analyses. Relative to the null distributions, GABA significantly decreased while glu/GABA ratio significantly increased (glutamate:tCr: p=0.097; GABA:tCr: p=0.015; glu/GABA ratio: p=0.009)% indicates p<0.05, ** indicates p<0.01.

As an additional control, we assessed changes in glu/GABA ratio during a subset of conditioning trials (Supplementary Fig. 9a) that were interleaved with the inference test trials during the MRI scan and shared the same temporal structure. Importantly, previous studies suggest performance on conditioning trials is not hippocampal-dependent^33^. During the conditioning trials, we observed no change in glu/GABA ratio during presentation of the visual cue or outcome, relative to the ITI period (Supplementary Fig. 9b–c).

### A hippocampal index for fluctuations in neocortical glu/GABA ratio

We next asked which brain regions coordinate this transient break in neocortical glu/GABA ratio during memory recall. The hippocampus is a promising candidate, given this brain region supports memory^3^ and shows activity modulation during the inference test (Fig. 2c). To test this possibility, we took advantage of our simultaneous fMRI-fMRS acquisition (Fig. 2a). We hypothesized that the increase in hippocampal BOLD signal observed during recall (Fig. 2c) should predict the increase in glu/GABA ratio observed in V1 (Fig. 3). In line with this prediction, across participants the hippocampal BOLD signal negatively predicted the relative concentration of GABA and positively predicted the increase in glu/GABA ratio in V1 (‘remembered’ versus ‘forgotten’ trials; Fig. 4a–b). Furthermore, across the imaged brain volume (Fig. 2a), only the hippocampus significantly predicted the increase in V1 glu/GABA ratio on ‘remembered’ versus ‘forgotten’ trials (Fig. 4c). Finally, this relationship between the hippocampus and glu/GABA ratio was specific to the recall period during the inference test (Fig. 4d, Supplementary Fig. 10).

**Figure 4.**
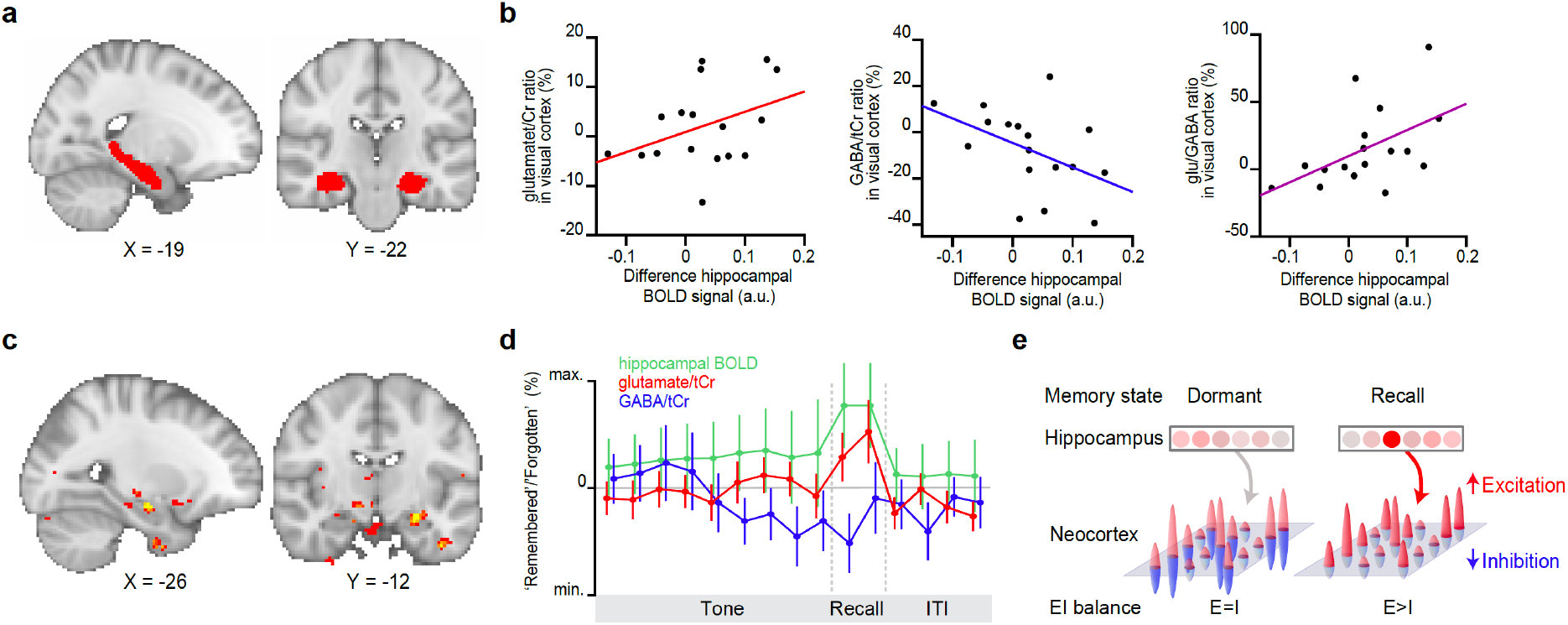
Hippocampal BOLD predicts neocortical glu/GABA ratio during recall. **a** Region of interest (ROI) in the hippocampus (red). **b** Across participants, the increase in hippocampal BOLD signal during ‘remembered’ compared to ‘forgotten’ trials positively predicted the decrease in GABA and the increase in glu/GABA ratio observed in V1 (Fig. 3b–d) (glutamate:tCr: r_16_=0.15, p=0.572; GABA:tCr: r_16_=−0.56, p=0.022; glu/GABA ratio: r_16_=0.52, p=0.033). **c** Across the imaged brain volume, for ‘remembered’ versus ‘forgotten’ trials the correlation between BOLD signal and V1 glu/GABA ratio was selectively observed in the left hippocampus (t16=11.25, p=0.005, whole-brain FWE corrected; Supplementary Table 6). **d** Moving average showing the ratio of ‘remembered’ to ‘forgotten’ trials during the inference test: hippocampal BOLD signal (green, n=19, range [−4:4]), glutamate:tCr (red, n=19, range [−8:8]), GABA:tCr (blue, n=19, range [−15:15]). Each point represents a 2.5s time bin (mean ± SEM). **e** Schematic illustrating how the hippocampus may facilitate memory recall of a sensory cue during a transient break in neocortical EI balance.

## DISCUSSION

The hippocampus is thought to provide an index for memories stored across distributed neocortical circuits^7–9^. However, the mechanism by which hippocampal activity is coordinated with neocortex to facilitate memory recall has remained unclear. Here, using time-resolved fMRI-fMRS in humans, we show that recall of a visual cue is accompanied by a dynamic increase in the ratio between glutamate and GABA in visual cortex. This transient increase in glu/GABA ratio in visual cortex is selectively predicted by activity in the hippocampus. Accordingly, we propose the hippocampus gates recall of memories stored across distributed neocortical circuits using a disinhibitory mechanism (Fig. 4e). This mechanism may explain how a memory index represented by the hippocampus selectively releases otherwise dormant representations stored across distributed neocortical circuits.

Memory recall via a disinhibitory mechanism may be supported by neural circuits identified in rodents, where glutamatergic projections from higher-order or interconnected brain regions have the capacity to instantiate highly specific disinhibition in cortical circuits^13,44,45^. For example, to enhance visual discrimination during attentional modulation, projections from the cingulate region of mouse frontal cortex modulate activity in V1 by targeting vasoactive intestinal polypeptide-expressing (VIP+) interneurons, which in turn preferentially target other interneuron subtypes to release excitatory principle cells from inhibitory control^13^. During memory recall, hippocampal projections may similarly instantiate highly specific disinhibitory control over cortical circuits to permit memory reinstatement. These findings are consistent with causal manipulations in humans showing that the hippocampus predicts memory expression in sensory neocortex unless neocortical glu/GABA ratio is disturbed^46^. Our results explain findings in humans showing that hippocampal GABA and glutamate can predict mnemonic control^47,48^ and may account for coordinated hippocampal-neocortical memory reinstatement reported in human imaging studies^37^ and intracranial recordings in epilepsy patients^10^. Moreover, hippocampal mediated neocortical disinhibition may potentially provide a signature for coordinated ripple-burst oscillatory activity between hippocampus and neocortex that has previously been observed in humans during memory recall^49^.

Our findings further speak to evidence reported from animal models showing that the ratio of excitatory to inhibitory synaptic conductance remains invariant, fluctuating around a stable set point^50^. While this may ensure that neurons and networks are neither hypo- nor hyper-excitable for prolonged periods, the exact E/I ratio is highly dynamic. Evidence in humans, animal models and theoretical models together suggest overall proportionality between excitation and inhibition is maintained to hold memories in a silent and dormant state^30–32,51^, thus protecting memories from interference caused by new learning^46,52^. Within this framework, memories must be released from inhibitory control to permit recall. While the precise mechanism may vary across brain systems and circuits, our data suggest disinhibition in V1 can release excitatory ensembles from balanced inhibition. Moreover, at the microcircuit level, disinhibition during memory recall has previously been identified following fear conditioning^53,54^. Thus, in addition to the established function of local disinhibition in promoting initial encoding of memory^11,45^, disinhibition may play a significant role in facilitating release and recall of previously learned but latent cortical associations.

During memory recall, we report a transient break in the glutamate/GABA ratio which can be attributed to a decrease in the concentration of MRS-derived GABA. The quality of the MRS data was comparable with other 7T MRS studies using unedited sequences to study glutamate and GABA in visual cortex^41,42,55,56^, as well as previous studies employing event-related fMRS^40,43,57^. While it is tempting to equate these changes in neurometabolite concentration with changes in synaptic activity, rapid changes in synaptic glutamate and GABA that accompany neurotransmitter release occur on a time-scale that is not possible to detect using the fMRI-fMRS sequence implemented here. Moreover, only a fraction of MRS-derived neurometabolite concentration reflects neurotransmitter release. MRS-derived measures fail to discriminate between different pools of glutamate and GABA (cytoplasmic, vesicular, or extracellular) and metabolites in different cellular compartments are maintained by a variety of different homeostatic mechanisms. MRS is considered most sensitive to unconstrained, intracellular metabolic pools that reside at relatively high concentration in the neuronal cytoplasm^58^. By comparison, changes in extracellular GABA of less than 100-fold are unlikely to be detectable using MRS^59^ and post-mortem studies suggest MRS is not sensitive to intracellular pools that reside in the mitochondria or vesicules^60,61^.

Interpretation of MRS-derived glutamate and GABA is further complicated by the fact that the release and recycling of glutamate and GABA constitute major metabolic pathways^21,22^. Yet, the metabolic and neurotransmitter pools are thought to be tightly coupled during anaesthesia, rest and certain stimulation protocols, with a 1:1 relationship reported between the rate of glutamine-glutamate cycling, which is necessary for glutamate and GABA synthesis, and neuronal oxidative glucose consumption, which indirectly supports neurotransmitter release among other processes^23–25^. Therefore, an increase in synaptic neurotransmission occurs together with an increase in synthesis of exogenous glutamate, which provides a precursor for GABA via the glutamate-glutamine cycle. During sensory stimulation a transient uncoupling has been observed with a short-lived mismatch between glucose utilization and oxygen consumption^62,63^, particularly during stimulation protocols that alternate between high intensity and quiescent periods^64^. Dynamic fluctuations in fMRS-derived glutamate and GABA reported here may therefore reflect transitions to new metabolic steady states^65^, which could reflect (if indirectly) relative shifts in EI equilibrium at the physiological level. During associative memory recall, the increase in glutamate/GABA ratio may be interpreted as an increase in synthesis of glutamate relative to degradation, with an opposing effect on GABA.

This interpretation is supported by a handful of previous studies showing event-related changes in MRS glutamate^40,43,57^ and GABA^66^, together with a growing body of evidence reporting a relationship between MRS-derived measures of neurometabolites and behaviour^67–69^. Nevertheless, it remains to be established whether unconstrained glutamatergic and GABAergic pools show event-related changes that are MRS-sensitive. To validate this interpretation of event-related fMRS it is important to leverage animal studies where more sensitive methods can be employed to relate fMRS measures to physiological parameters. Here, by implementing an inference task in VR, we operationalize memory recall using the exact same paradigm previously employed in rodents^33^. Therefore, in addition to engaging attention and memory-dependent inference, “opening the box” to find a reward in the VR environment approximated the process of rodents finding a reward from a dispenser in a 3D environment. By using VR, the findings presented here may be compared to data acquired in animal models in ongoing future research. In this manner, VR paradigms in humans may provide a basis from which to gain insight into the cellular and circuit mechanisms that underlie macroscopic measures of excitation and inhibition. This may prove particularly useful for establishing a more detailed understanding of the relationship between fMRS-derived measures of glutamate and GABA and physiological measures of EI balance.

Previous fMRS protocols typically employ a ‘block’ design, where a static measure of the concentration of glutamate and GABA is achieved by averaging the spectra across a time-window that may span several minutes. The clear limitation of this approach is that dynamic changes in glutamate and GABA are not assessed in relation to cognitive processes and ongoing behaviour that occur on a much faster scale. With the increase in availability of ultra-high field MRI scanners and the development of more advanced sequences^70^, fMRS has emerged as a viable method to detect dynamic changes in neurochemicals in both healthy and clinical populations^65^. Although there are currently only a handful of event-related fMRS studies, together with our data, these suggest fMRS is highly sensitive to detecting task-relevant dynamic changes in glutamate and GABA^71^. For example, in the lateral occipital complex fMRS demonstrates differences in glutamate in response to presentation of objects versus abstract stimuli^57^, and in the left anterior insula fMRS reveals a transient increase in glutamate with exposure to painful stimuli^40^. fMRS-derived glutamate is even sufficiently sensitive to detect repetition suppression effects in the lateral occipital complex^43^, mirroring analogous effects reported in fMRI^72^. Here, we further illustrate that within a 3 second window delineated by the question period in the inference task, the temporal resolution of fMRS is sufficient to relate transient changes in glutamate and GABA to memory performance. fMRS therefore provides a promising tool to capture real-time, task-relevant changes in neurometabolites, on a time scale equivalent to task-based fMRI. Assessing whether the temporal resolution of fMRS can be further improved will likely prove an important step in refining fMRS in the future.

During associative memory recall, the transient increase in glu/GABA ratio reported in our data can primarily be accounted for by a significant decrease in the concentration of MRS-derived GABA, which was in turn predicted by the hippocampal BOLD signal. Notably, detecting dynamic changes in GABA is challenging for two key reasons: the concentration of GABA in human brain tissue is relatively low and the spectral peaks for GABA overlap with other, more abundant neurochemicals^73–75^. While the most common approach to detecting MRS-derived GABA involves using a J-difference spectral editing technique to separate peaks that derive from GABA from overlapping peaks^76,77^, here we use a non-edited sequence (sLASER). While spectral editing may provide higher precision^56^, this occurs at the cost of a larger volume of interest and longer TEs, which makes it less suitable for event-related fMRS^78,79^. Moreover, direct comparisons between edited and non-edited sequences at 7T reveal no significant difference in the concentration of GABA measurements^56^. Therefore, together with studies reporting dynamic changes in GABA with sensory stimulation^55,80^, our data illustrates how a non-edited sequence can provide sufficient data quality for measuring dynamic changes in MRS-derived GABA, which cannot be explained by changes in compounds at higher-concentration that have overlapping peaks (i.e. glutamate or NAA, Supplementary Fig. 8). Moreover, compared to spectral editing, our approach comes with the advantage of simultaneously measuring dynamic changes in GABA and glutamate, together with 17 other neurometabolites.

Disturbances in EI balance are thought to underlie a number of neuropsychiatric conditions, including schizophrenia, autism, epilepsy and Tourette’s syndrome^65,81,82^. While previous studies report inconsistencies in MRS-derived measures of glutamate and GABA in these patient populations, this may be attributed to differences in brain region, cognitive state and imaging protocol, among other factors. Here, by using both fMRS and fMRI to reveal a signature change in glu/GABA ratio that relates to hippocampal BOLD signal, behavioural performance and cognition, our findings present a potential target for clinical investigation.

In summary, using time-resolved fMRI-fMRS we report a transient increase in glu/GABA ratio in V1 during associative recall of a visual cue. This increase in glu/GABA ratio can be attributed to a decrease in the concentration of MRS-derived GABA, which is predicted by activity in the hippocampus. By unveiling this coordination between the hippocampus and neocortex, we show how the hippocampus may have the capacity to selectively modulate and disinhibit memories represented in neocortex. This mechanism may explain how the hippocampus plays a key role in memory recall, by indexing the release of specific memories stored across distributed neocortical circuits.

## METHODS

### Participants

22 healthy human volunteers were included in the study (mean age of 22.8 ± 0.74 years, 4 males). All experiments were approved by the University of Oxford ethics committee (reference number R43594/RE001). All participants gave informed written consent. For one participant, we were unable to collect combined fMRI-fMRS data due to time constraints during scanning. Two participants were excluded from the fMRI and fMRS analyses due to technical difficulties which resulted in the auditory cues not being fully audible during the inference test. Notably, there was no significant effect of sex on either behavioural performance or MRS measures of glu/GABA ratio during the inference test (Supplementary Table 1).

### Virtual reality environment

The virtual reality (VR) environment was coded using Unity 5.5.4f1 software (Unity Technologies, CA United States). The environment was designed to simulate an open field environment previously used to investigate memory and inference in mice33. By incorporating VR, our experimental design is therefore suitable for making cross-species comparisons in the future. This may prove important when seeking to establish a more refined interpretation of fMRS in relation to neural circuit mechanisms.

The environment included a square-walled room with no roof (Fig. 1b). To help evoke the experience of 3D space and aid orientation within the VR environment, each wall of the environment was distinguished by colour (dark green, light green, dark grey or light grey), illumination (two walls were illuminated while the other two were in shadow) and by the presence of permanent visual cues. The permanent visual cues included clouds in the sky, a vertical black stripe in the middle of the light green wall, a horizontal black strip across the light grey wall, and a wooden box situated in one corner of the environment. A first-person perspective was implemented and participants could control their movement through the virtual space using the keyboard arrows (2D translational motion) and the mouse-pad (head tilt). Movement through the environment elicited the sound of footsteps. Within the VR environment participants were exposed to a range of different sensory stimuli, in accordance with the three-stage inference task described below.

### Three-stage inference task

In the VR environment (Fig. 1b) humans performed an inference task (Fig. 1a). The rationale for using an inference task to assess mechanisms responsible for associative memory was three-fold. First, evidence in both humans and mice shows that performance on this inference task requires associative memory recall33. Second, in mice, inference, but not first-order associative recall, is hippocampal dependent^33–35^, thus providing an opportunity to investigate hippocampal dependent associative memory recall. Third, the task can be deployed across humans and rodents, which may allow future investigation of the cellular mechanisms that underlie non-invasive measures reported here.

The task was adapted from associative inference and sensory preconditioning tasks described elsewhere^33,83,84^ and involved 3 stages performed across 3 consecutive days, respectively (Fig. 1a,c). The first and second stages were performed outside the scanner while the third stage was performed inside the scanner (Fig. 1c). At the start of the experiment the pairings between auditory, visual and outcome cues were randomly assigned for each participant.

On day 1, participants performed the ‘observational learning’ stage (Fig. 1a), during which participants were required to learn at least 40 (out of 80 total) auditory-visual associations via mere exposure. In total, there were 4 visual cues, each associated with 20 different auditory cues. Auditory cues constituted 80 different complex sounds (e.g. natural sounds or those produced by musical instruments) that were played over headphones. Visual cues constituted 4 different unique patterned panels which could appear on the walls of the environment (Fig 1a,b,e). To control for potential spatial confounds, two of the visual cues were always presented on the same wall, the assignment of which was randomized for each participant. The two remaining visual cues were ‘nomadic’, meaning that with each presentation they were randomly assigned to one of the four walls.

Training during the observational learning stage occurred within the VR environment and was divided into 8 sub-sessions. In each sub-session, participants controlled their movement within the VR environment and were presented with 20 trials in which 10 different auditory-visual associations, different in each sub-session, were each presented twice, in a random order. On each trial an auditory and visual cue were presented serially and contiguously: 8 s auditory cue followed by 8 s of the associated visual cue, followed by an ITI of 5 s (Supplementary Fig. 1a). Participants were given the choice to repeat the sub-session if they so wished. After the sub-session, learning of auditory-visual associations was monitored outside the VR environment, using an observational learning test coded in Matlab 2016b using Psychtoolbox (version 3.0.13). On each trial of the observational learning test, 1 auditory cue from the sub-session was presented, followed by presentation of 4 different visual cues (Supplementary Fig. 1b). Participants were instructed to select the visual cue associated with the auditory cue using a button press response within 3 s, and only at the end of the test were participants given feedback on their average performance. Each auditory cue in the sub-session was presented 2 times. Participants were required to repeat training in the VR environment (including the observational learning test) until they obtained at least 50% accuracy for auditory-visual associations in the sub-session (chance level: 25%).

After obtaining at least 50% accuracy on the observational learning test for each sub-session, participants were given an ‘overview’ memory test (Supplementary Fig. 1b). The memory test had the same format as the observational learning test used for each sub-session, except that it included all 80 auditory cues, each of which was presented 3 times. Training on the observational learning stage was terminated when participants reached >50% accuracy on this ‘overview’ memory test (Supplementary Fig. 1e). If participants failed to reach >50% accuracy, training in the VR environment was repeated for those sub-sessions with poor performance. Those participants that failed to reach >50% accuracy on the ‘overview’ memory test (n=3) did not proceed to day 2 and were thus not included in the experiment.

On day 2, participants performed the ‘conditioning’ stage (Fig. 1a), during which they learned that two of the four visual cues (set 1) predicted delivery of a rewarding outcome (virtual silver coin, as above) on 80% of trials, while the other two visual cues (set 2) predicted delivery of a neutral outcome (virtual wood-chip, as above) on 100% of trials. The outcomes were delivered to a wooden box situated in the corner of the environment. To harvest the value of a virtual silver coin (monetary reward later converted to 20 pence per coin) or a virtual woodchip (no value, 0 pence), participants were required to first collide with the wooden box, which caused its walls to disappear, before colliding with the coin or wood-chip which was accompanied by a ‘collision’ sound. The outcome cues were only available for 10 s. The cumulative total value of harvested reward was displayed in the upper left corner of the computer screen.

Training during the conditioning stage occurred within the VR environment and on each trial, participants were presented with a visual cue and outcome which were presented serially and contiguously: visual cue (8 s) followed by outcome delivery to a wooden box (available for 6 s) (Supplementary Fig. 1c). Participants were instructed to only look in the wooden box after the visual cue was presented and instructed to leave the wooden box before the next trial. The inter-trial interval (ITI) was 2 s.

Learning during the VR conditioning training was monitored using a conditioning test coded in Matlab 2016b using Psychtoolbox (version 3.0.13). On each trial of the conditioning test, participants were presented with a still image of a visual cue before being asked to indicate the probability of reward using a number line (Supplementary Fig. 1d). Participants were given 3 s to respond and were only given feedback on their average performance at the end of the test. Participants were required to repeat the VR conditioning training and conditioning test until they performed the test with 100% accuracy (Supplementary Fig. 1f).

Finally, on day 3, participants first repeated the conditioning test. Participants then entered the 7T MRI scanner and performed the ‘inference test’ (Fig. 1a, c–d), together with a subset of conditioning trials (Supplementary Fig. 9a) (see *fMRI-fMRS scan task* below). Immediately after exiting the scanner, participants were given a surprise associative memory test to assess which auditory-visual associations they remembered and which they had forgotten (Fig. 1e). The memory test was equivalent to the test performed on day 1 during the observational learning (Supplementary Fig. 1b), with 3 trials for each auditory stimulus. Performance on auditory-visual associations was categorised as correct if participants scored 3/3 for that auditory cue on the subsequent surprise memory test. Performance on auditory-visual associations was categorised as incorrect if participants scored 0/3 or 1/3 for that auditory cue on the subsequent surprise memory test (i.e. no different from chance). Trials where participants scored 2/3 were not categorised as either correct or incorrect due to their ambiguity. The behavioural performance measured on the post-scan associative memory test (Fig. 1f) was a more sensitive measure of memory accuracy than behavioural performance measured during the inference test, with a lower chance level (associative memory test: 4 choice options with 25% chance level; inference test: 2 options with 50% chance level) and more repeats of each auditory cue (associative memory test: 3 repeats; inference test: 1 repeat). For this reason, performance on the inference test during the scan was assessed post-hoc using performance from both the inference test and the post-scan associative memory test (see *Trial categorisation during the inference test*, Fig. 2b).

### fMRI-fMRS scan task

The inference test was incorporated into the fMRI-fMRS scan task. This provided an opportunity to measure neural responses to associative memory recall required for inferential judgements. The scan task included two different trial types: inference test trials (Fig. 1d) and conditioning trials (Supplementary Fig. 9a). For both types of trial participants viewed a short video taken from the VR training environment. The videos were presented via a computer monitor and projected onto a screen inside the scanner bore. On each trial the duration of the video was determined using a truncated gamma distribution with mean of 7 s, minimum of 4 s and maximum of 14 s. During the inference test trials, the video of the VR environment was accompanied by an auditory cue, played over MR compatible headphones (S14 inset earphones, Sensimetrics). Visual cues were not displayed during these trials: the auditory cues were presented in isolation. At the end of the video, participants were presented with a question asking: ‘Would you like to look in the box?’, with the options ‘yes’ or ‘no’ (Fig. 1d). Participants were required to make a response within 3 s using an MR compatible button box and their right index or middle fingers. No feedback was given. To infer the appropriate outcome participants were instructed to use the learned structure of the task. The inference test thus provided an opportunity to investigate memory recall: to infer the correct choice participants needed to recall the appropriate visual cue associated with the auditory cue (Fig. 1g). Conditioning trials were interleaved with inference test trials to minimise extinction effects. During conditioning trials, the video of the VR environment orientated towards a visual stimulus displayed on one of the four walls (Supplementary Fig. 9a). At the end of the video, participants were presented with a still image of the associated outcome for that visual cue (Supplementary Fig. 9a). After each trial (inference or conditioning) a cross was presented in the centre of the screen during an inter-trial interval of varying length, determined using a truncated gamma distribution (mean of 2.7 s, minimum of 1.4 s, maximum of 10 s).

To control for potential confounding effects of space, each video during the inference test involved a trajectory constrained to a 1/16 quadrant of the VR environment, evenly distributed across the different auditory cues. Across conditioning trials, each visual cue was presented 16 times, once in each possible spatial quadrant. The fMRI-fMRS scan task was evenly divided across 2 scan blocks, each of which lasted 15 minutes. The fMRI-fMRS scan task was then repeated (2 more scan blocks) using a higher quality multiband fMRI sequence (not reported here).

### fMRI-fMRS data acquisition

The fMRI-fMRS scan task was performed inside a 7 Tesla Magnetom MRI scanner (Siemens) using a 1-channel transmit and a 32-channel receive phased-array head coil (Nova Medical Inc., USA) at the Wellcome Centre for Integrative Neuroimaging Centre (University of Oxford). Current 7T radio-frequency (RF) coil designs suffer from B1^+^ inhomogeneity. To overcome this, we positioned two 110 × 110 × 5 mm^3^ Barium Titanate dielectric pads (4:1 ratio of BaTiO3:D2O, relative permittivity around 300) over occipital lobe, causing a “hotspot” in the proximal B1^+^ distribution at the expense of distal regions^85^. For each participant, a T1-weighted structural image was acquired to inform placement of the MRS voxel in visual cortex, and to correct for geometric distortions and perform co-registration between EPIs, consisting of 176 0.7 mm axial slices, in-plane resolution of 0.7 × 0.7 mm^2^, TR = 2.2 s, TE = 2.96 ms, and field of view = 224 mm. For each participant, a field map with dual echo-time images was also acquired (TE1 = 4.08 ms, TE2 = 5.1 ms, whole-brain coverage, voxel size 2 × 2 × 2 mm^3^).

Fig. 2a shows a diagram of the combined fMRI-fMRS sequence, based on a sequence developed by Hess et al.^86^, and previously used to compare the BOLD signal in V1 with measures of glutamate36. In the same TR of 4s, BOLD-fMRI (3D EPI, resolution 2.3 × 2.3 × 2.2 mm^3^; flip angle=5°, repetition time TR_epi_= 59 ms, TE=29 ms, field of view 200 mm, 32 slices) and fMRS data (2 × 2 × 2 cm^3^ voxel positioned in the occipital lobe, centered along the midline and the calcarine sulcus) were acquired. fMRS data were acquired using short-echo-time semi-localisation by adiabatic selective refocusing (semi-LASER) pulse sequence (TE=36 ms, TR_mrs_=4 s) with VAPOR water suppression and outer volume suppression87. A delay between fMRI and fMRS acquisition (250 ms) was inserted to minimize potential eddy current effects from the EPI read-out86. Compared to an uncombined contemporary MR sequences (e.g. multiband EPI and semi-LASER MRS), the fMRS was of comparable quality, while the quality of the fMRI component was compromised. On average, 457 fMRS spectra were acquired over the two scanning blocks (SD: 35.62).

In addition to the fMRI-fMRS sequence acquisition, an additional set of fMRI data (reported elsewhere33 and not shown here) was acquired using a multiband EPI sequence (50 1.5 mm thick transverse slices with 1.5 mm gap, in-plane resolution of 1.5 × 1.5 mm^2^, TR=1.512 s, TE= 20 ms, flip angle = 85°, field of view 192 mm, and multi-band acceleration factor of 2). To increase SNR in brain regions for which we had prior hypotheses, both the fMRI sequences were restricted to partial brain coverage (Fig. 2a, covering the occipital and temporal lobes) to shorten the EPI TR, thus acquiring more measurements.

### Trial categorisation during the inference test

Trials during the inference test were categorised into two conditions, ‘remembered’ and ‘forgotten’ (Fig. 2b). To obtain the most accurate estimate of associative memory recall during the inference test our definition for ‘remembered’ and ‘forgotten’ derived from behavioural performance on both the inference test and the post-scan associative memory test. Trials where participants made both the correct inference during the inference test *and* subsequently remembered the auditory-visual association during the post-scan associative memory test were classified as ‘remembered’. Trials where participants made *either* the incorrect inference during the inference test *or* subsequently forgot the auditory-visual associations during the post-scan associative memory test were classified as ‘forgotten’.

### fMRS metabolite quantification and analysis

For each scan run, fMRS data from 19 subjects was preprocessed separately in MRspa, a semi-automated MATLAB routine (https://www.cmrr.umn.edu/downloads/mrspa/). The unsuppressed water signal acquired from the same VOI was used to remove residual eddy current effects and combine individual coil spectra. Spectra were corrected for frequency and phase variations induced by participants’ motion, and the residual water component was removed using Hankel Lanczos Singular Value Decomposition (HLSVD). For each participant, spectra from all blocks were frequency aligned to account for frequency differences between blocks.

Spectra were then analysed in an event-related manner. For each participant, the preprocessed spectra were first assigned to the tone/question/ITI periods by aligning the time stamps for the spectra to the time stamps for each event recorded during the inference task. Then, spectra acquired within the tone/question/ITI periods were selected for analysis. Next, these selected spectra were separated into two categories according to task performance, ‘remembered’ or ‘forgotten’ (Fig. 2b, see *Trial categorisation during the inference test*), before being analysed using LCModel. Participants (n=1) with less than 8 spectra for either the ‘remembered’ or ‘forgotten’ conditions were excluded from the fMRS analysis, as previous studies report minimal change in test-retest CoVs when going from 8 to 16 spectra88. Metabolite concentrations for the average ‘remembered’ and the average ‘forgotten’ spectrum were quantified in turn using LCModel89 within the chemical shift range 0.5 to 4.2 ppm. The concentration of each metabolite was assessed relative to the concentration of total Creatine (Creatine + phosphocreatine, tCr), thus providing effective control for variation in voxel tissue and cerebral spinal fluid (CSF) in the fMRS voxel used across participants. A basis set containing stimulated model spectra of alanine (Ala), aspartate (Asp), ascorbate/vitamin C (Asc), glycerophosphocholine (GPC), phosphocholine (PCho), creatine (Cr), phosphocreatine (PCr), GABA, glucose (Glc), glutamine (Gln), glutamate (Glu), glutathione (GSH), myo-inositol (myo-Ins), Lactate, N-acetylaspartate (NAA), N-acetylaspartylglutamate (NAAG), phosphoethanolamine (PE), scyllo-inositol (scyllo-Ins), taurine (Tau) and experimentally measured macromolecules was used. To evaluate the dynamic range of metabolites between ‘remembered’ and ‘forgotten’ conditions, it was not appropriate to use the default settings in LCModel that normalise metabolite estimates such as GABA to constrain the dynamic range. We therefore removed these prior constraints within LCModel by setting the ‘*nratio*’ parameter to 0. Estimates normalised to tCr were multiplied by 8, as per convention.

Changes in the relative concentration of glutamate and GABA between ‘remembered’ and ‘forgotten’ conditions were evaluated together with ‘glu/GABA ratio’ which we defined as the ratio of glutamate to GABA39. We defined the change in glutamate, GABA and glu/GABA for ‘remembered’ vs ‘forgotten’ trials as a ratio, as follows:

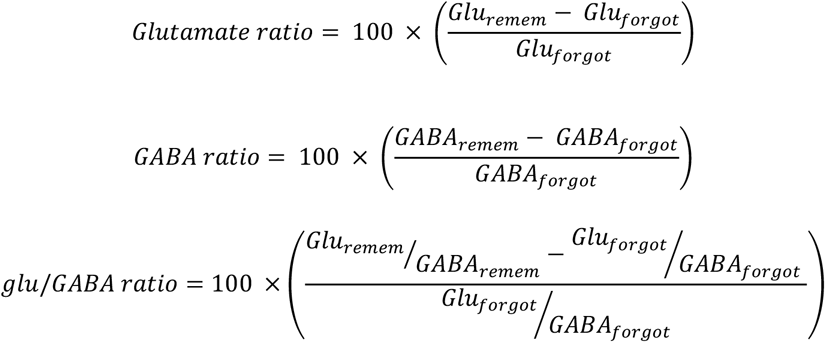

Where Glu and GABA represent the ratio of glutamate and GABA to total Creatine, respectively, during the tone/question/ITI period of ‘remembered’ or ‘forgotten’ trials. This ratio effectively controls for variation in voxel tissue and CSF fraction in the MRS voxel used across participants.

Further, to control for differences in the number of ‘remembered’ and ‘forgotten’ spectra, we compared the group mean difference between ‘remembered’ and ‘forgotten’ trials against a null distribution generated by permuting the trial labels while preserving differences in number of trials for each participant. On each of 5000 permutations, the condition labels (‘remembered’, ‘forgotten’) were shuffled for each participant using MATLAB’s random number generator. The relative metabolite concentrations for each condition were then estimated in LCModel and the difference between conditions computed. The group mean for each permutation was then added to the null distribution. The difference between ‘remembered’ and ‘forgotten’ conditions derived from the unshuffled data was then compared against the null distribution generated from the shuffled data (Fig. 3d–f; Supplementary Fig. 5–8).

### fMRI preprocessing and GLMs

Preprocessing of MRI data was carried out using SPM12 (http://www.fil.ion.ucl.ac.uk/spm/). First, the anterior commissure was set to the origin in the anatomical images and in the first volume of each fMRI block, with equivalent transformations applied to all other images within the same block. Second, to account for magnetic field inhomogeneities, images were corrected for signal bias, realigned to the first volume, corrected for distortion using field maps, normalised to a standard EPI template. To remove low frequency noise from the pre-processed data, a high-pass filter was applied to the data using SPM12’s default settings. For each participant and for each scanning block, the resulting fMRI data was analysed in an event-related manner using a general linear model (GLM). The GLM was applied to data from both scan task blocks. In addition to the explanatory variables (EVs) of interest (described below), 6 additional scan-to-scan motion parameters produced during realignment were included in the GLM as nuisance regressors to account for motion-related artefacts in each task block. The output of the first-level analysis was then smoothed using a 5-mm full-width at half maximum Gaussian kernel before being entered into a second level analysis. The sensitivity of our analysis pipeline to detecting stimulus evoked BOLD activity patterns benefitted from applying the first-level GLM to unsmoothed data and only including smoothing prior to the second level analysis (Supplementary Fig. 2). One participant was excluded from the fMRI analyses as the quality of fMRI data in the fMRI-fMRS sequence was too poor to ensure reliable pre-processing.

For the first-level analyses, three different GLMs were used. Each GLM included 15 EVs per block. In the first GLM, the first 8 EVs accounted for the question period in the inference test, divided according to performance of the subject (‘remembered’ or ‘forgotten’, see *Trial categorisation during the inference test*), before being further divided according to the 4 possible visual cues to which the auditory cues were associated. The next 4 EVs accounted for presentation of the visual cue during the video of all conditioning trials, divided according to the 4 different visual cues. The final 3 EVs accounted for presentation of the auditory cue during the video in all inference test trials, the question period in all remaining inference test trials (i.e. trials not categorized as ‘remembered’ or ‘forgotten’), and the presentation of the outcome in all conditioning trials. To decorrelate the EVs modelling the auditory and visual cues from those EVs modelling the question and outcome, respectively, the duration of events within EVs modelling the auditory and visual cues was set using a box-car function to 4 s, i.e. the minimum duration of the video. The duration of events within EVs modelling the question/outcome were set to the duration of the question/outcome. All EVs were then convolved with the hemodynamic response function.

In the second and third GLMs, the same EVs were included, however the first 8 EVs accounted for the auditory cue period in the inference test (second GLM), or the inter-trial interval in the inference test (third GLM). In both cases, the EVs were divided according to performance of the subject (‘remembered’ or ‘forgotten’), as in the first GLM.

### Univariate fMRI analysis and statistics

Using the output of the GLMs we assessed the difference in the univariate BOLD response between ‘remembered’ and ‘forgotten’ trials during the inference test (as defined in Fig. 2b, *Trial categorisation during the inference test*). The contrast of interest therefore involved contrasting EVs [1:4] (‘remembered’) with EVs [5:8] (‘forgotten’), using the first GLM (see above). The resulting contrast images (‘remembered’-‘forgotten’) for all participants were entered into a second-level random effects ‘group’ analysis. We set the cluster-defining threshold to p<0.01 uncorrected before using whole-brain family wise error (FWE) to correct for multiple comparisons, with the significance level defined as p<0.05 (Fig. 2c, Supplementary Table 3).

### Assessing the relationship between fMRI and fMRS

To assess the relationship between event-related hippocampal BOLD signal and event-related fMRS measures from V1, we used an anatomical ROI for the hippocampus (Fig. 4a). Capitalising on variance across participants, the relationship between the BOLD signal for ‘remembered’-‘forgotten’ within this ROI was compared with equivalent changes in glutamate, GABA and glu/GABA ratio using a Spearman rank correlation. To assess the selectivity of these effects to the recall period (question) during the inference test, control analyses were performed using the output of the second and third GLMs, together with equivalent measures of glutamate, GABA and glu/GABA ratio (Supplementary Fig. 10).

Next, to assess the relationship between fMRS and the BOLD signal across the entire imaged brain volume, we repeated the second-level random effects ‘group’ analysis using the output of the first GLM, but now included group-level covariates for the change in glutamate and GABA for ‘remembered’-‘forgotten’ (i.e. Fig. 3a), along with 2 ‘nuisance’ regressors that accounted for unwanted variance attributed to differences in age and sex. To identify brain regions where the BOLD signal for ‘remembered’:‘forgotten’ predicted changes in glu/GABA ratio, we contrasted the explanatory variables on the covariates for glutamate and GABA (glutamate – GABA) to generate a single contrast to test statistical significance. We set the cluster-defining threshold to p<0.01 uncorrected before using whole-brain family wise error (FWE) to correct for multiple comparisons, with the significance level defined as p<0.05 (Fig. 4c, Supplementary Table 6).

To visualize the time course of fMRI and fMRS across the inference test trials, we estimated a moving average for both datasets, where each time bin constituted a 2.5 s time window shifted by 0.5 s in each iteration (Fig. 3c, Fig. 4d, Supplementary Fig. 10c–d). To account for the jitter in the length of the video and in the ITI, trials that stopped short were excluded from analyses for that time bin. Thus, to ensure each time bin contained a similar number of spectra, those time bins at the tail end of the jitter (final 3 time bins during the video and the final 2 time bins of the ITI) were enlarged to include broader time windows. For the fMRS, for each participant the ‘remembered’ and ‘forgotten’ spectrum were calculated for each time bin, and the ratio estimated to give a measure of ‘remembered’:‘forgotten’ for both glutamate and GABA (Fig. 3c, Fig. 4d, Supplementary Fig. 10c–d).

For the fMRI, for each participant, and for each time bin during the inference test trial, the time course of the preprocessed BOLD signal was extracted from the hippocampal ROI (Fig. 4a) and from two control ROIs defined using a 12 mm sphere within our partial epi volume (Fig. 2a). The first control region was positioned at the junction between parietal and occipital cortex (‘parietal-occipital cortex’) while the second control region was positioned within the brainstem (Supplementary Fig. 10c–d). For each ROI, the obtained signal for each trial was resampled using a resolution of 400 ms and regressed against an explanatory variable indicating those trials that were remembered. To control for differences in baseline BOLD at the start of the trial, we also included a ‘nuisance’ explanatory variable indicating whether the previous trial was ‘remembered’. We then plotted the normalized averaged fMRI regression coefficient for ‘remembered’ vs ‘forgotten’ together with the equivalent glu/GABA ratio time course (Fig. 4d; Supplementary Fig. 10c–d).

## DATA AVAILABILITY

Upon publication, data for all figures will be made available made available on GitHub (https://github.com/rskool/memory_recall).

## CODE AVAILABILITY

Upon publication, code used to analyse the data for all figures will be made available made available on GitHub (https://github.com/rskool/memory_recall).

## Acknowledgements

We would like to thank Aaron Hess for advice regarding the combined fMRI-fMRS sequence. R.S.K. is supported by an EPSRC/MRC-funded studentship (EP/L016052/1). A.S. is supported by a Wellcome Trust studentship (203836/Z/16/Z). D.D. is supported by the Biotechnology and Biological Sciences Research Council UK (BBSRC UK award BB/N0059TX/1) and the MRC (Programme MC_UU_12024/3). H.C.B. is supported by the John Fell Oxford University Press Research Fund (Grant 153/046), a seed grant from the Wellcome Centre for Integrative Neuroimaging, a Junior Research Fellowship from Merton College (University of Oxford) and the Medical Research Council (MRC) UK (MC_UU_12024/3). The Wellcome Centre for Integrative Neuroimaging is supported by core funding from the Wellcome Trust (203139/Z/16/Z).

## Author contributions

All of the authors contributed to the preparation of the manuscript. R.S.K., A.S., D.D., and H.C.B. designed the study; U.E.E. and I.B.I. developed the combined fMRI-fMRS sequence; R.S.K., A.S., U.E.E., and H.C.B. acquired the data; R.S.K., A.S., U.E.E., W.T.C., and H.C.B. analysed the data.

## Competing interests

Authors declare no competing interests.

## Materials & Correspondence

Correspondence and requests for materials should be addressed to.

## Supplementary Information

### Supplementary Figures

**Supplementary Figure 1.**
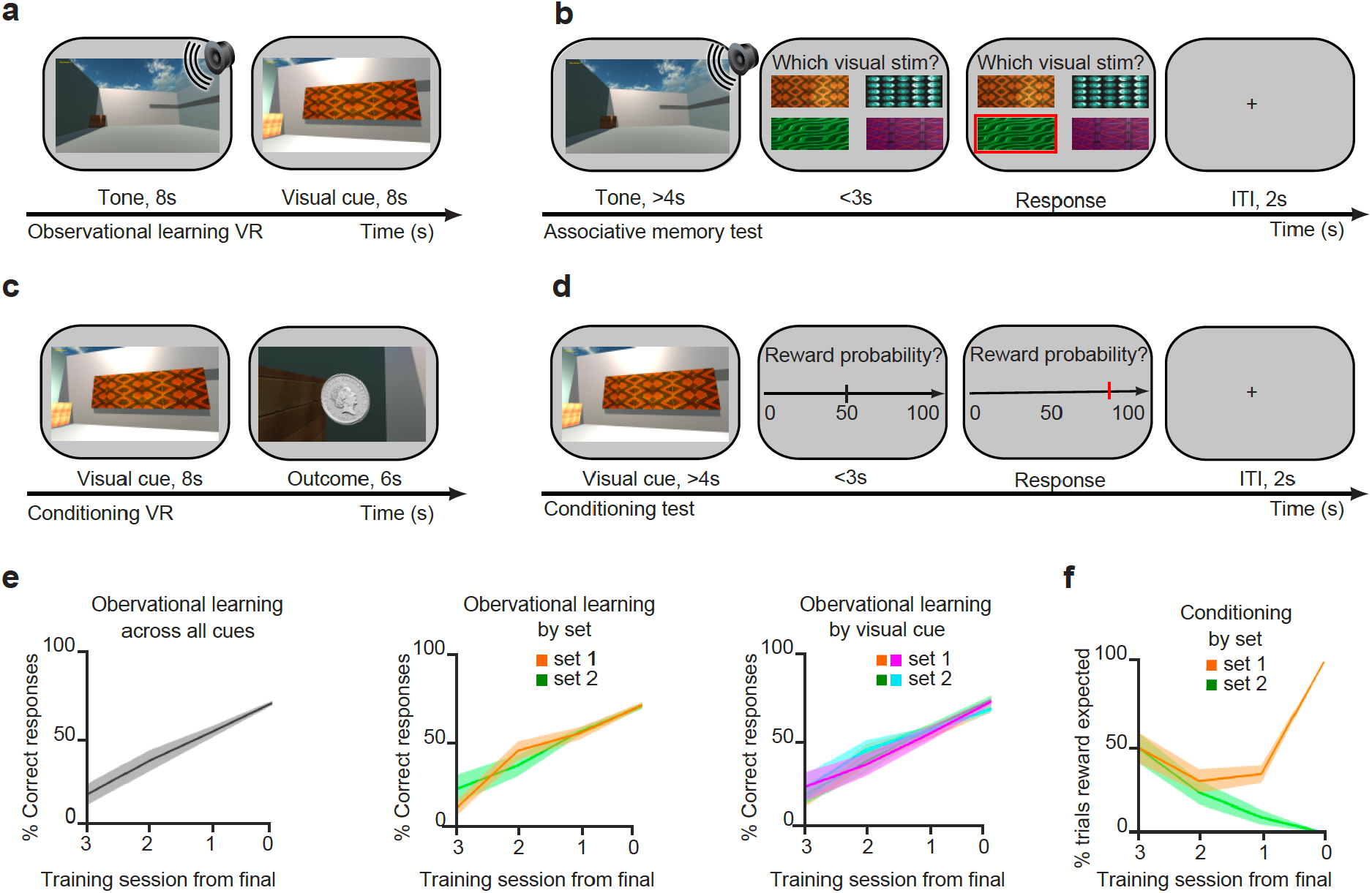
Behavioural training and performance. **a-b** On day 1, during the ‘observational learning’ stage, participants learned to associate each of the 80 different auditory cues with one of 4 possible visual cues. **a** The observational learning stage was performed in a VR environment (Fig. 1b). **b** Learning was monitored using an associative memory test, in the absence of feedback. **c-d** On day 2, during the ‘conditioning’ stage, participants learned to associate each of the four visual cues with one of two possible outcomes (monetary reward for set 1; neutral woodchip for set 2) (Fig. 1a). **c** The conditioning stage was performed in a VR environment (Fig. 1b). **d** Learning was monitored using a conditioning test, in the absence of feedback. **e** Left: Participants performed the observational learning task until they showed recall accuracy on the associative memory test of at least 50% of the 80 possible auditory-visual pairs. Middle: On the associative memory test, used to monitor performance during the observational learning, there was no difference in accuracy between auditory-visual pairs in set 1 (rewarded) and set 2 (neutral). Right: On the associative memory test there was no difference in accuracy between auditory cues associated with the four different visual cues. **f** On day 2, participants performed the conditioning task until they reached 100% accuracy on all visual-outcome associations in the conditioning test.

**Supplementary Figure 2.**
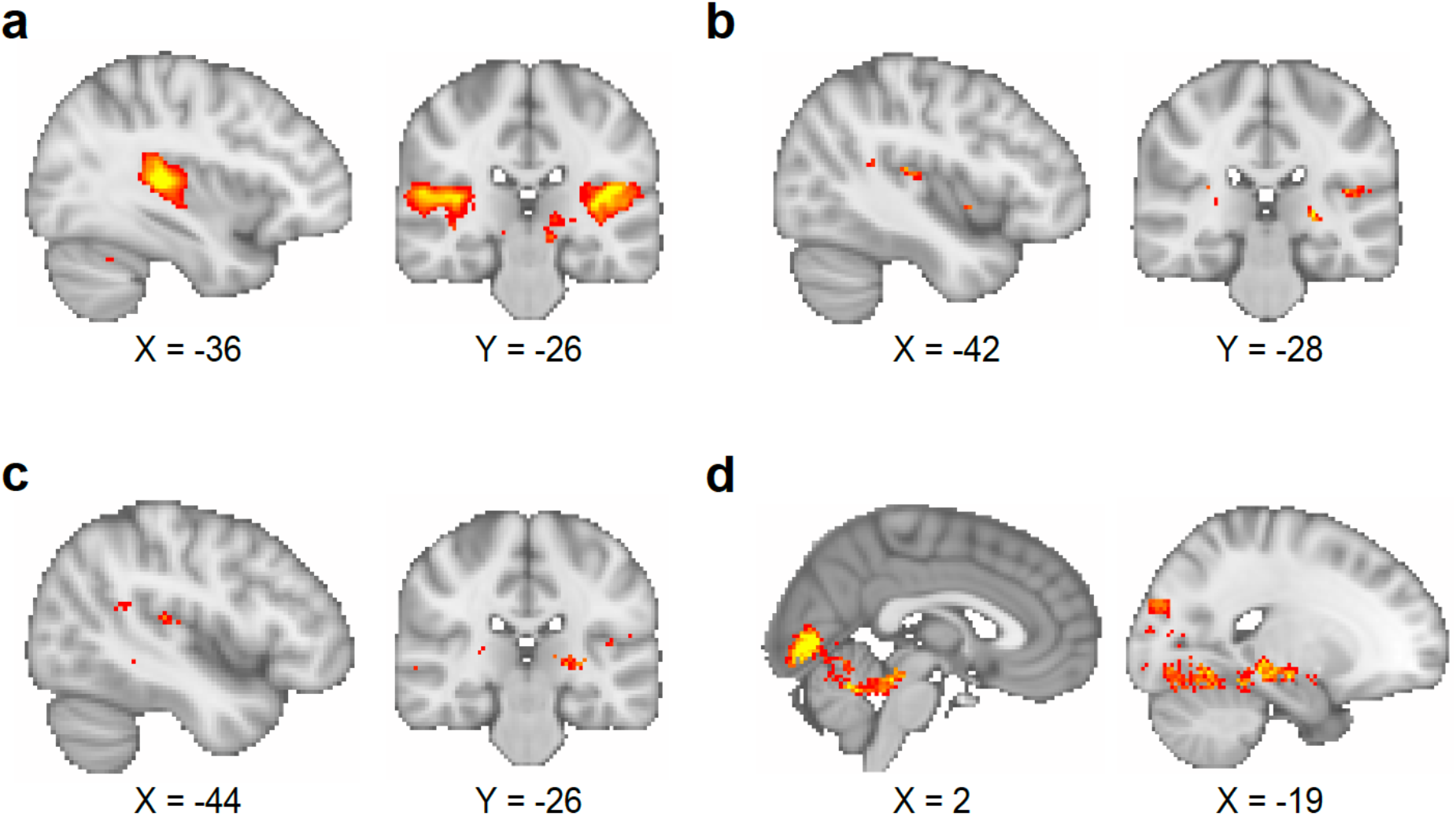
Comparison of different smoothing parameters applied to fMRI data. **a-c** The BOLD signal response to auditory cues in the inference test was used to assess the effect of different smoothing parameters (contrast of interest: [all trials during inference test –all trials during conditioning]). As noted in the *Methods*, the quality of the fMRI data in the interleaved fMRI-fMRS sequence was compromised relative to contemporary standards for 7T fMRI. The smoothing parameters applied to the data influenced the reliability of the analysis, as illustrated here. **a** Smoothing at the second-level using a 5 mm kernel, approximately two times the voxel size, as recommended90, gave significant BOLD signal in bilateral auditory cortex (left auditory cortex, t_17_=8.94, p<0.001; right auditory cortex, t_17_=6.76, p<0.001; whole-brain FWE corrected; thresholded at p<0.001 uncorrected for visualisation purposes only). This smoothing protocol was used for all analyses presented in the main figures. **b** Smoothing at the first-level using a 5 mm kernel, did not give significant BOLD signal in auditory cortex (whole-brain FWE corrected; thresholded at p<0.01 uncorrected for visualisation purposes only). **c** Smoothing at the first-level using an 8 mm kernel (default for SPM) did not give significant BOLD signal in auditory cortex (whole-brain FWE corrected; thresholded at p<0.01 uncorrected for visualisation purposes only). **d** When contrasting ‘remembered’ and ‘forgotten’ trials in the inference test, smoothing at the first-level using an 8 mm kernel (as in ****c****) gave similar results to those presented in Fig. 2c, despite the absence of a main effect response to auditory cues as shown in ****c****. Thus, as shown in Fig. 2c, during the question period in the inference test trials (Fig. 1a, d), BOLD signal in the visual cortex and the hippocampus was significantly higher for ‘remembered’ versus ‘forgotten’ auditory cues (‘remembered’ –‘forgotten’, visual cortex: t_17_=6.01, p<0.001; right hippocampus: t_17_=7.58, p<0.001; whole-volume FWE-corrected; thresholded at p<0.01 uncorrected for visualisation purposes only).

**Supplementary Figure 3.**
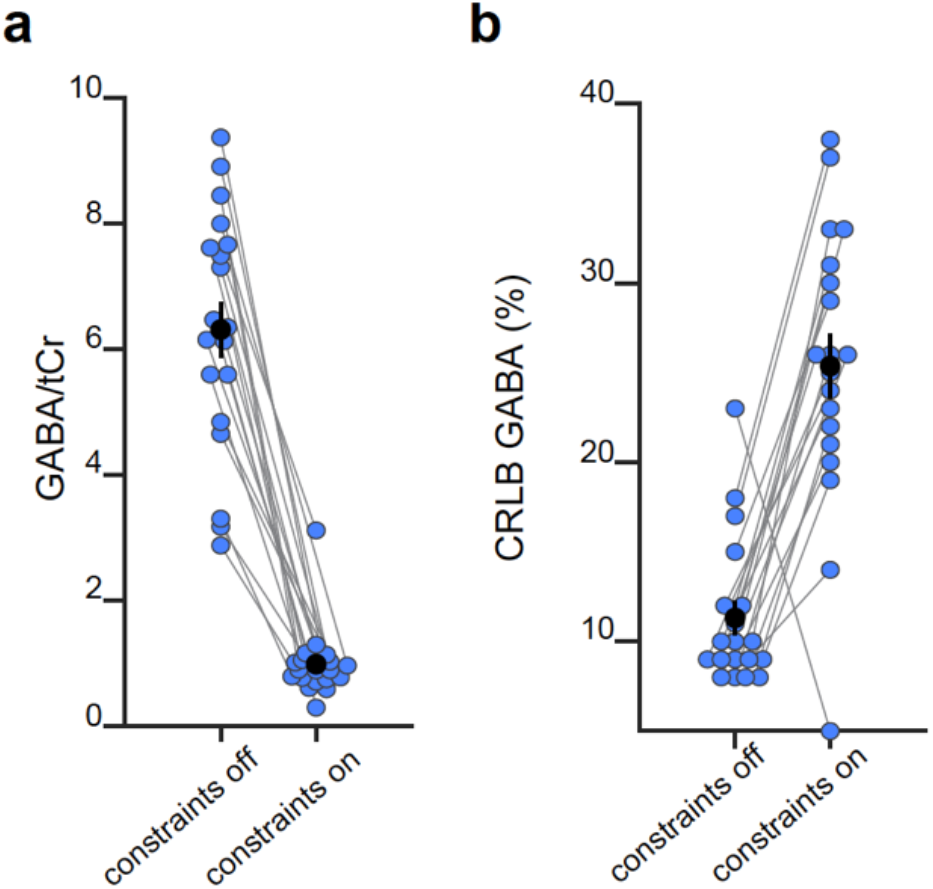
Comparison of GABA estimates and uncertainty in the model fit with and without model constraints. **a** When using a model fit that constrains metabolite values within a predefined (‘physiologically plausible’) range (i.e. ‘constraints on’), GABA values are reduced together with their dynamic range. However, the assumptions of these constraints are not suitable for detection of dynamic fluctuations in glu/GABA ratio. **b** When using a model fit that constrains the metabolite values within a predefined range, the uncertainty of the GABA estimates increase, as indicated by the Cramér–Rao Lower Bounds (CRLBs). This shows that the reliability of the model fit for GABA is higher when model constraints are removed, namely the approach required to measure dynamic fluctuations in glu/GABA ratio.

**Supplementary Figure 4.**
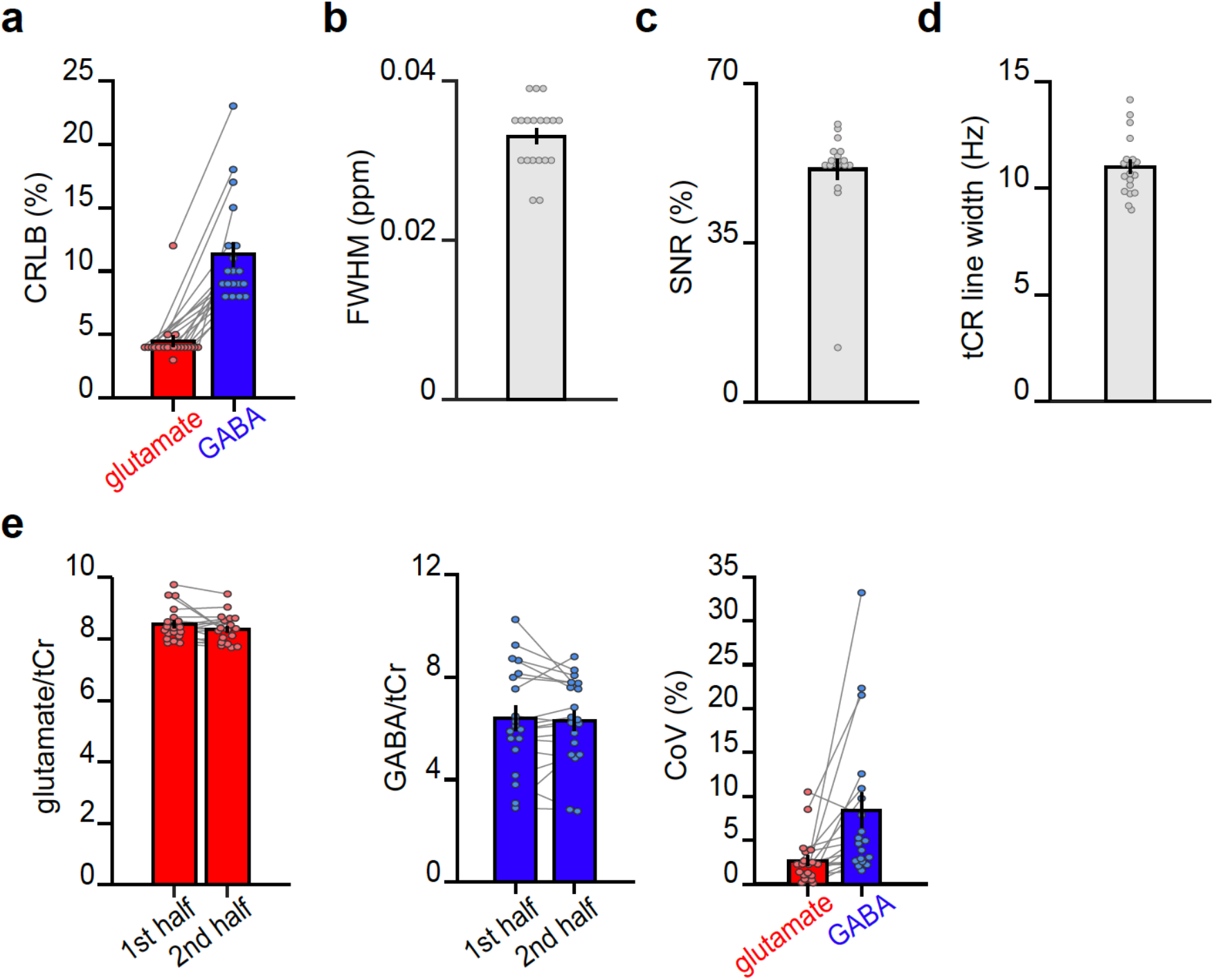
MRS data quality metrics across all spectra. For each subject, quality metrics across all MRS spectra were assessed (mean total number of spectra: 457). **a** The average Cramér-Rao Lower Bound (CRLB) for glutamate was 4.47±0.42% and the average CRLB for GABA was 11.31±0.94%. (mean ± SEM) **b** The average full width at half max (FWHM) as determined by LCModel was 0.033±0.001ppm (mean ± SEM). **c** The average Signal to Noise Ratio (SNR) as determined by LCModel was 51.1± 2.37%. (mean ± SEM). **d** The average line width of the total Creatine (tCr) peak was estimated to be 11.01±0.32Hz. (mean ± SEM). **e** The intra-subject coefficient of variance (CoV) was estimated by splitting the dataset into two equal halves (on average 278 spectra in each half) and analysing each half in LCModel. We defined CoV as the standard deviation between the 2 halves divided by their mean. The intra-subject CoV for glutamate was 2.68±0.62%, and the intra-subject CoV for GABA was 8.41±1.96% (mean ± SEM). These findings demonstrate stability in our MRS measurements over the course of the scan task. Notably, this analysis differs from standard estimates of intra-subject CoV where test-retest is assessed across two separate scanning sessions.

**Supplementary Figure 5.**
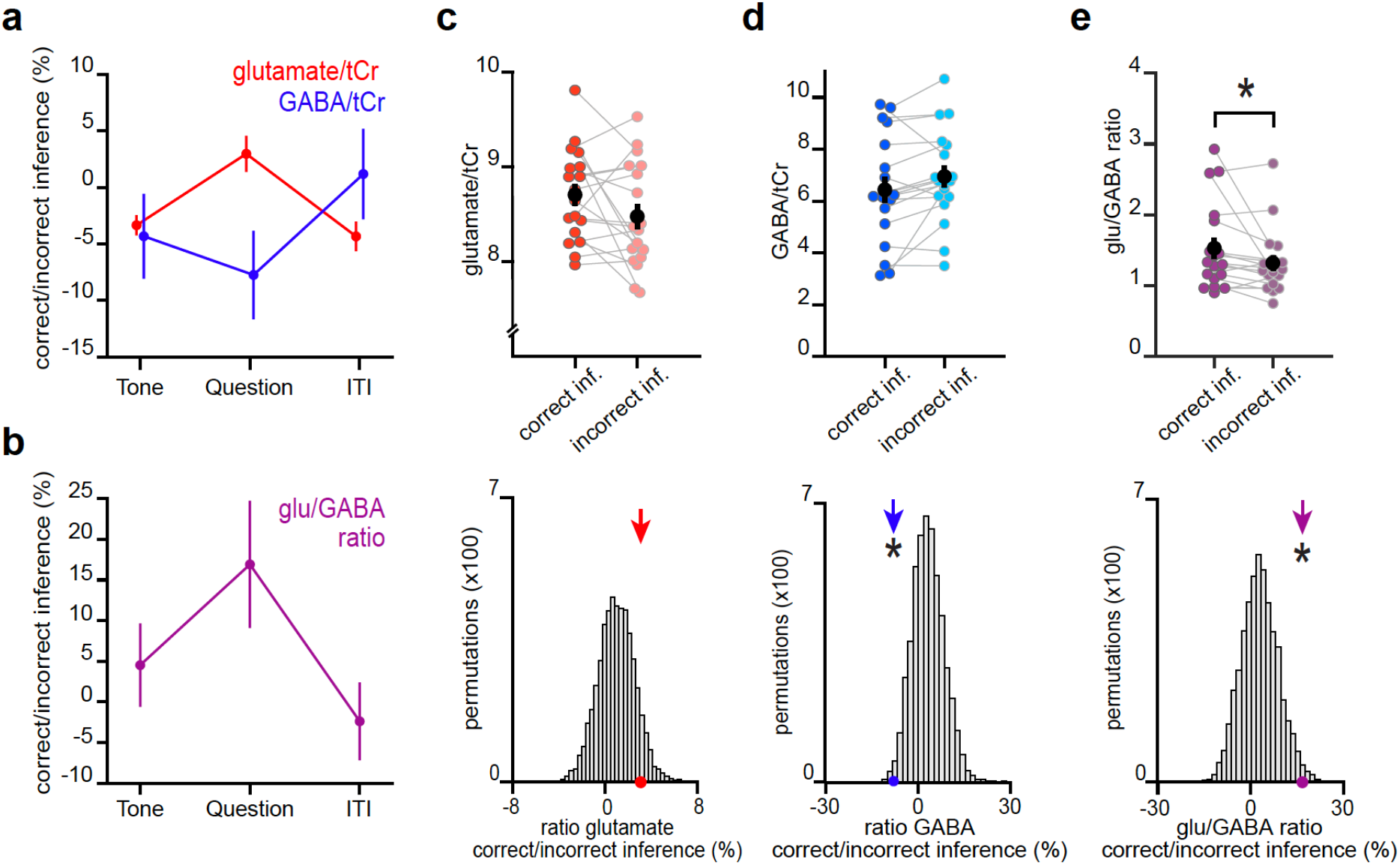
An increase in glu/GABA ratio in V1 during memory recall are also observed when categorising trials into ‘remembered’ and ‘forgotten’ using a less conservative approach. Here we repeated the analyses in Fig. 3 using only performance on the inference test to categorize trials from the inference test into ‘remembered’ (correct inference) and ‘forgotten’ (incorrect inference). **a-b** Similar to using the more conservative definition for ‘remembered’ and ‘forgotten’ shown in Fig. 3, glu/GABA ratio significantly increased during ‘remembered’ versus ‘forgotten’ trials (‘correct inference’:‘incorrect inference’, glu/GABA ratio: t_17_=2.16, p=0.045). This break in glu/GABA ratio was not observed during the ‘tone’ (~7 s) or ‘ITI’ (~2.7 s) periods (‘Tone’, glu/GABA ratio: t_18_=0.88, p=0.391; ‘ITI’, glu/GABA ratio: t_18_=−0.50, p=0.623). **c-e** Upper row: Metabolite values and glu/GABA ratio during the question period for ‘remembered’ and ‘forgotten’ trials (mean ± SEM). Lower row: Comparison of the mean ratio of ‘correct inference’ to ‘incorrect inference’ (coloured arrows) against null distributions generated by permuting the trial labels (x5000 to control for any potential biases in the analyses. Relative to the null distributions, GABA significantly decreased while glu/GABA ratio significantly increased (glutamate:tCr: p=0.088; GABA:tCr: p=0.017; glu/GABA ratio: p=0.011). * indicates p<0.05.

**Supplementary Figure 6.**
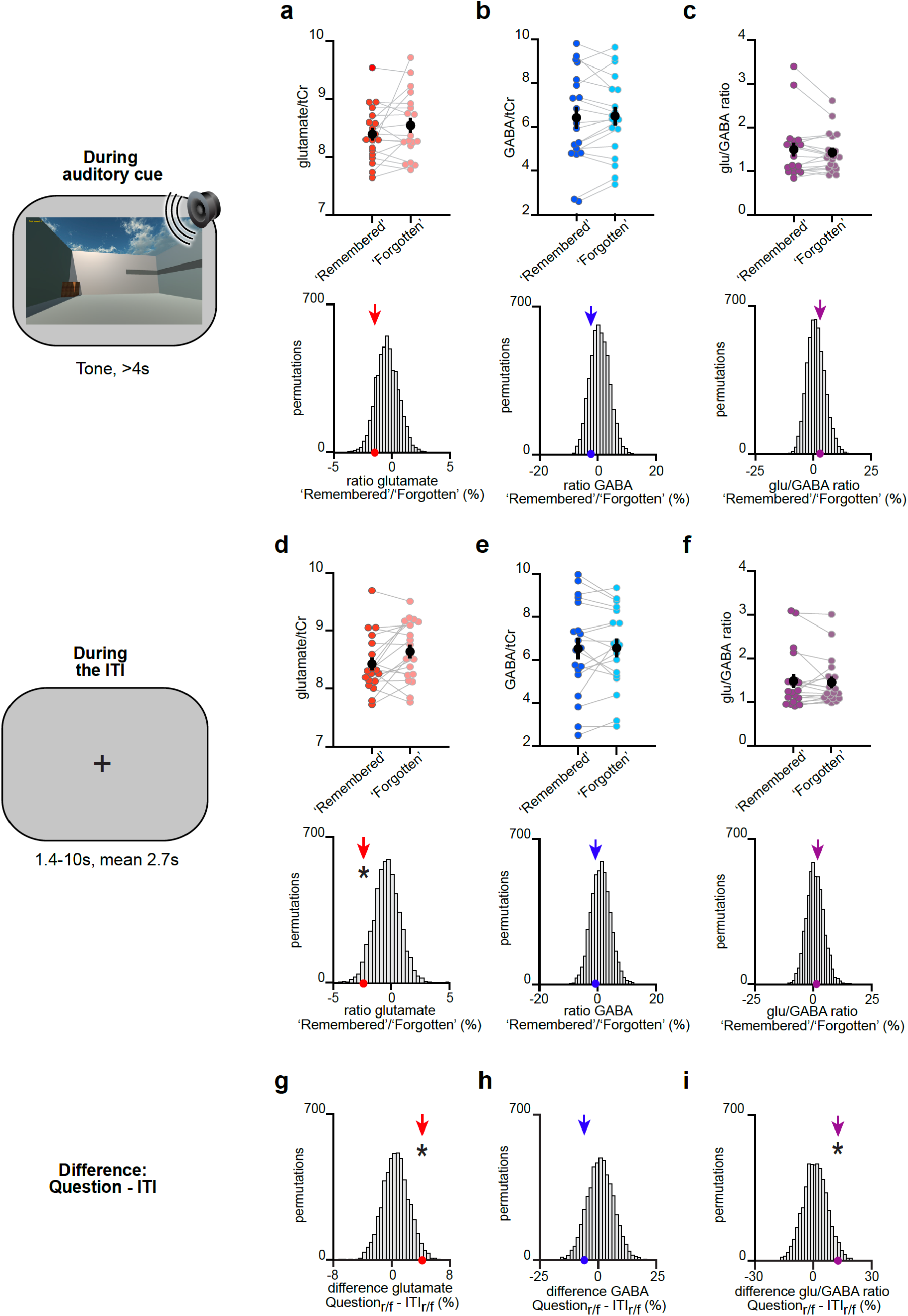
The change in glu/GABA ratio is transient and only observed during memory recall. **a-f** Upper rows: The difference in metabolite ratios for ‘remembered’ versus ‘forgotten’ trials during the auditory cue period (a-c) and ITI period (d-f) (mean ± SEM). Lower rows: To control for any biases due to differences in the number of ‘remembered’ and ‘forgotten’ trials, we compared the group mean (coloured arrows) against a null distribution generated by permuting the trial labels (x5000). MRS voxel shown in Fig. 2d. **a-c** During the auditory cue period, there was no significant increase in glu/GABA ratio between ‘remembered’ and ‘forgotten’ trials (‘remembered’:‘forgotten’, glutamate:tCr: t_18_=1.40, p=0.180; GABA:tCr: t_18_=0.80, p=0.433; glu/GABA ratio: t_18_=0.74, p=0.468). Similarly, there was no significant difference between any of the group means and their respective null distributions (glutamate:tCr: p=0.107; GABA:tCr: p=0.191; glu/GABA ratio: p=0.314). **d-f** During the ITI period, there was no significant increase in glu/GABA ratio between ‘remembered’ and ‘forgotten’ trials (‘remembered’:‘forgotten’, glutamate:tCr: t_18_=−2.27, p=0.040; GABA:tCr: t_18_=0.31, p=0.761; glu/GABA ratio: t_18_=0.31, p=0.766). For GABA and glu/GABA ratio, there were no differences between any of the group means and their respective null distributions, although a significant decrease was observed for glutamate (glutamate:tCr: p=0.029; GABA:tCr: p=0.271; glu/GABA ratio: p=0.438). **g-i** To test whether the break in glu/GABA ratio was transient and only observed during memory recall, we compared our measure of glu/GABA ratio for ‘remembered’:‘forgotten’ during the question period (‘Question’) to the period immediately after (‘ITI’) on the same trial. To control for the difference in numbers of trials between conditions, we compared the difference of the group means to a permuted null distribution. Compared to the respective null distributions, a significant difference was observed for glutamate and glu/GABA ratio between the group means for ‘Question’ versus ‘ITI’ (glutamate:tCr: p=0.021; GABA:tCr: p=0.110; glu/GABA ratio: p=0.034). ‘r/f’ indicates ‘remembered’:‘forgotten’,*indicates p<0.05.

**Supplementary Figure 7.**
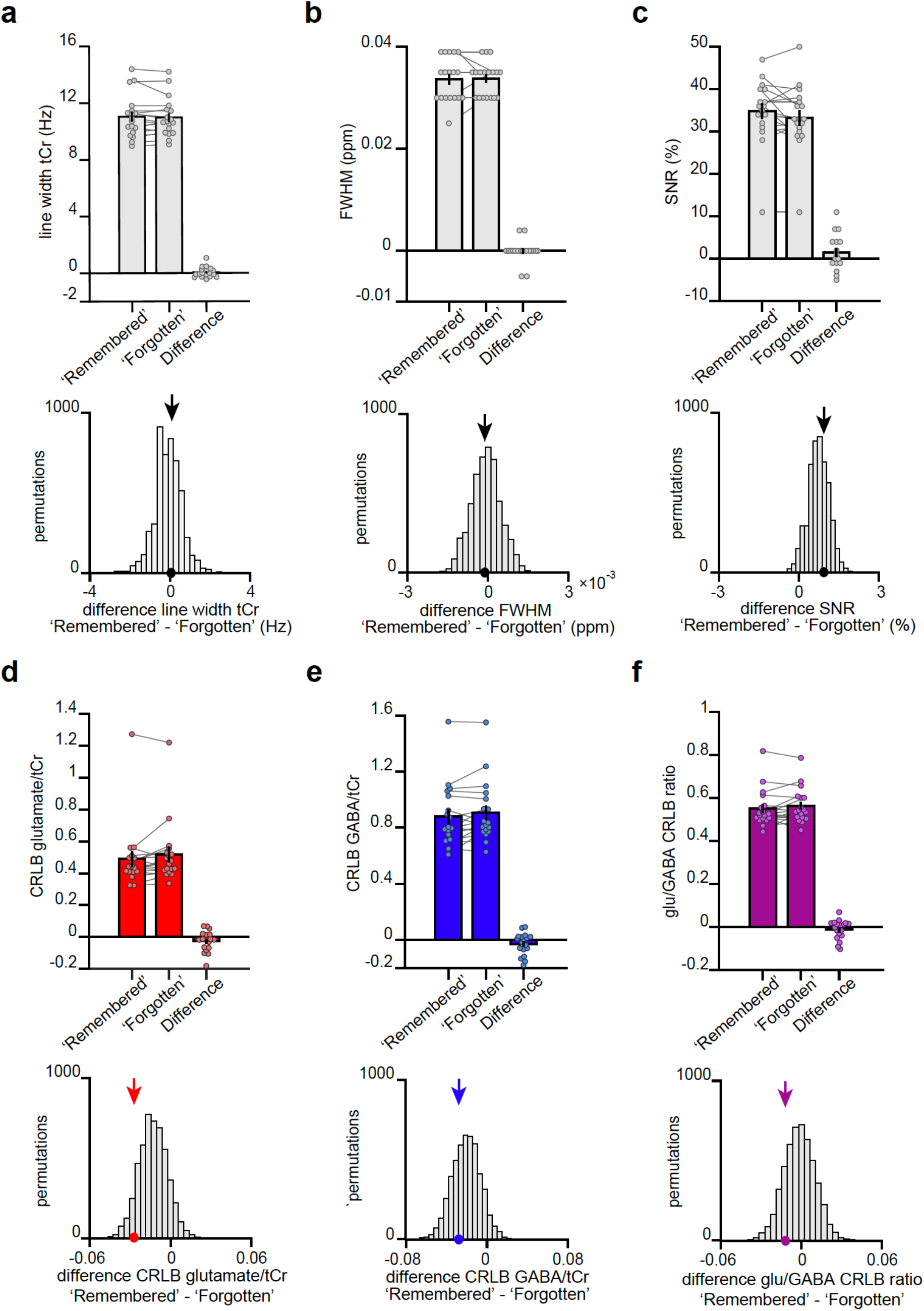
The transient break in glu/GABA ratio observed during recall cannot be explained by changes in data quality metrics or goodness of model fit. **a-b** At 7T, increases in BOLD effects alter T2* of metabolite signals, a phenomenon that results in line narrowing of all signals in the spectrum91. This phenomenon is most discernible on the strongest singlets such as total creatine (tCr). To assess the reliability of our fMRS measures we therefore quantified the difference in line width between our conditions of interest. These analyses show no significant difference in line width between categories ‘remembered’ and ‘forgotten’. Thus, as linewidth was matched across categories there was no evidence for a category-specific bias in BOLD-related confounds for metabolite values reported in Fig. 3. **a** The line width of the tCr peak was estimated for each participant for both the ‘remembered’ and ‘forgotten’ spectra acquired during the question period of inference test. Across participants we observed no difference in tCr line width between our two conditions of interest (‘remembered’ –‘forgotten’: t_17_=0.61, p=0.552) (mean ± SEM). To control for systematic differences in the number of ‘remembered’ and ‘forgotten’ trials we compared the group mean difference in tCr (black arrow) against a null distribution generated by permuting the trial labels (x5000) and re-estimating line width difference for ‘remembered’ - ‘forgotten’. Again, there was no significant difference between the group mean difference in tCr and the null distribution (p=0.403). **b** To further quantify differences in line width for each participant’s ‘remembered’ and ‘forgotten’ spectra, we compared the full width at half maximum (FWHM). Between our two conditions of interest we observed no significant difference in FWHM (‘remembered’ –‘forgotten’: t_17_=0.21, p=0.832) (mean ± SEM). There was no significant difference between the FWHM group mean for ‘remembered’ –‘forgotten’ and its null distribution generated as described in (*a*) (p=0.467). **c** To verify that the differences between ‘remembered’ and ‘forgotten’ cannot be attributed to a difference in signal strength, we assessed the SNR for both conditions. We observed no significant difference in SNR (‘remembered’ - ‘forgotten’: t_17_=1.55, p=0.140). In addition, there was no significant difference between the SNR group mean difference for ‘remembered’ –‘forgotten’ and its null distribution (p=0.312). The positive shift of the null distribution can be explained by the difference in number of trials between the ‘remembered’ and ‘forgotten’ conditions (Supplementary Table 5). **d-f** To verify that the differences observed between ‘remembered’ and ‘forgotten’ spectra cannot be attributed to differences in model fit, we assessed the CRLB for our metabolites of interest. We observed no significant difference in CRLBs (‘remembered’ - ‘forgotten’, glutamate: t_17_=−1.73, p=0.101; GABA: t_17_=−1.53, p=0.145; glu/GABA ratio: t_17_=−1.13, p=0.276). In addition, there was no significant difference between the CRLB group mean differences for ‘remembered’ –‘forgotten’ and their respective null distributions (glutamate: p=0.084; GABA: p=0.301; glu/GABA ratio: p=0.183). The negative shift of the glutamate and GABA CRLB null distributions is related to the difference in SNR between ‘remembered’ and ‘forgotten’; CRLB is lower for conditions with higher SNR, indicating a more confident model fit.

**Supplementary Figure 8.**
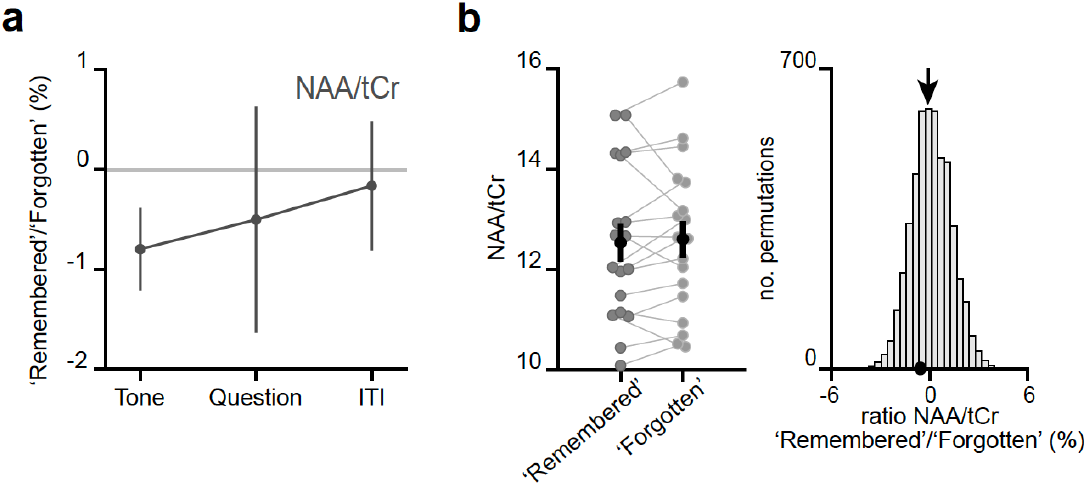
The changes in metabolite concentrations cannot be attributed to changes in NAA:tCr. **A** No significant change between ‘remembered’ and ‘forgotten’ trials was observed for NAA during the ‘Question’ period of inference trials (remembered:forgotten NAA:tCr: t_17_=−0.44, p=0.663). Notably, NAA has overlapping peaks with GABA but is found at higher concentration. **b** Left: Metabolite values during the question period for ‘remembered’ and ‘forgotten’ trials (mean ± SEM). Right: Ratios between ‘remembered’:’forgotten’ (black arrows) against null distributions generated by permuting the trial labels. Relative to the null distributions, the mean NAA ratio showed no significant difference (NAA:tCr: p=0.259).

**Supplementary Figure 9.**
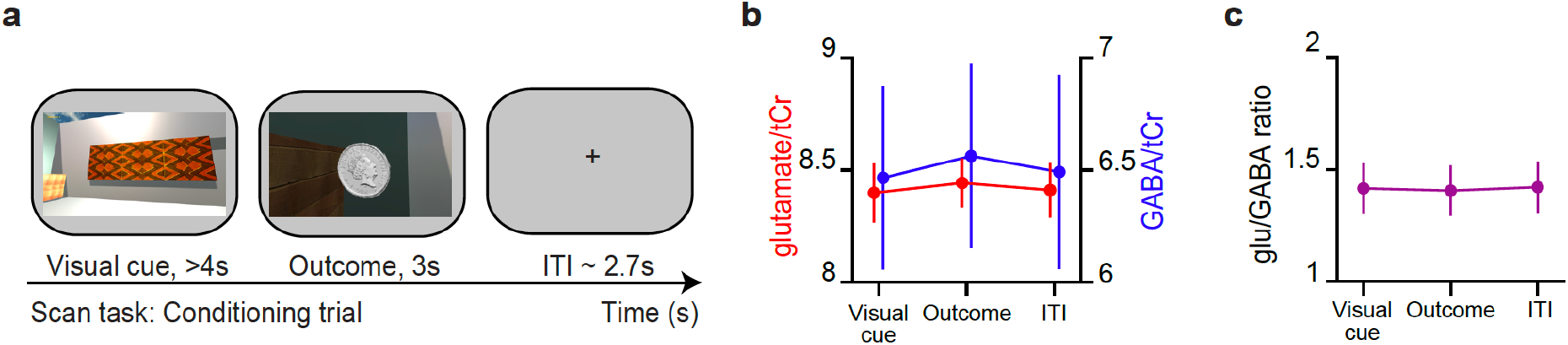
During conditioning trials, no difference in glu/GABA ratio was observed. **a** Example conditioning trial encountered during the MRI scan task. **b** The concentration of glutamate:tCr and GABA:tCr during the ‘visual cue’, ‘outcome’ and ‘ITI’ periods of conditioning trials. **c** glu/GABA ratio did not change during presentation of the visual cue or outcome, relative to the ITI period (‘Visual cue’-’ITI’: t_18_=−0.11, p=0.915; ‘Outcome’ - ‘ITI’: t_18_=−0.21, p=0.833).

**Supplementary Figure 10.**
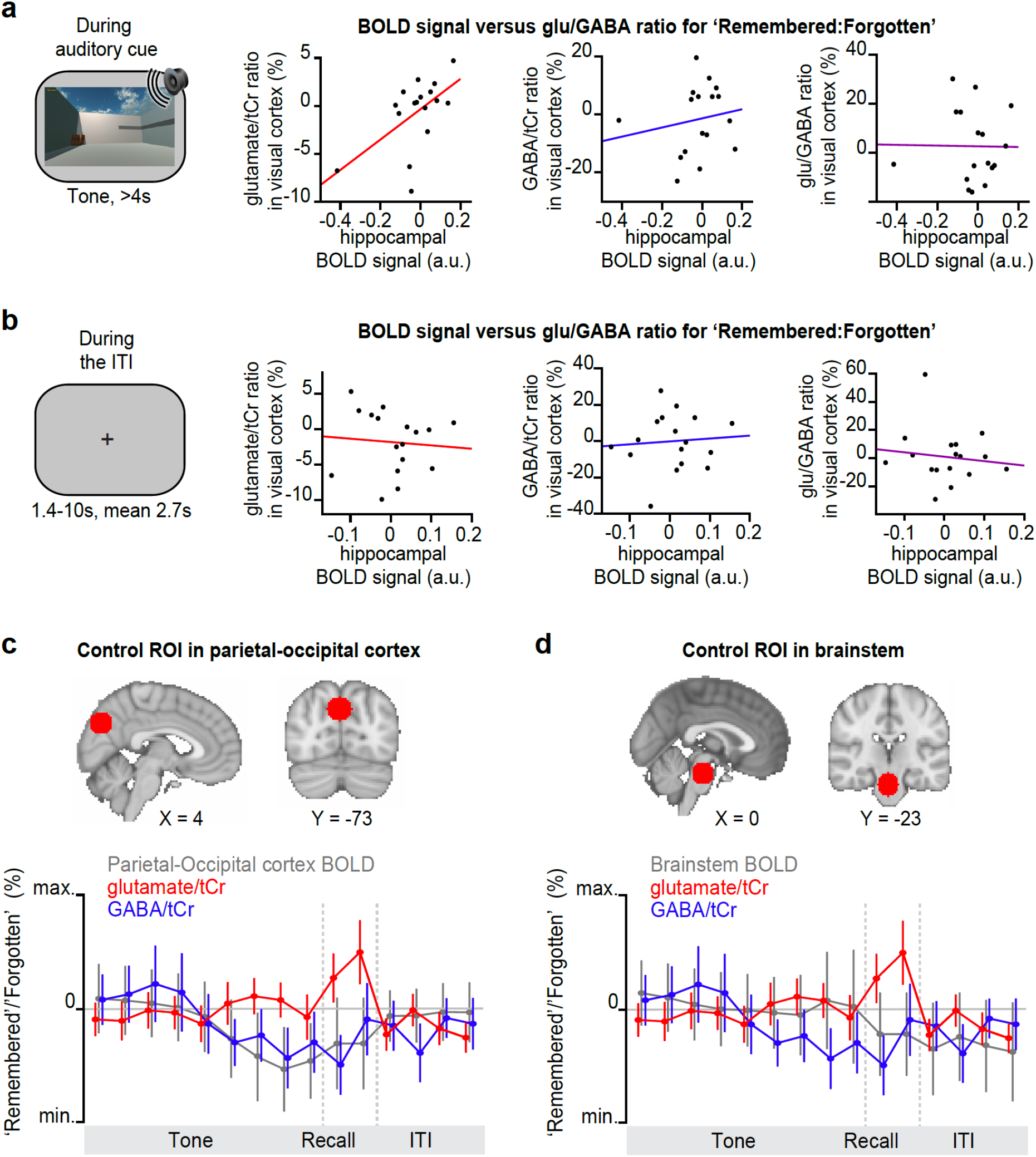
Before and after memory recall, the hippocampal BOLD signal did not positively predict glu/GABA ratio in V1. **a-b** During ‘remembered’ relative to ‘forgotten’ trials on the inference test (the question period, Fig. 1e) we observed a significantly positive correlation between the hippocampal BOLD signal and glu/GABA ratio in visual cortex (Fig. 4b–c). To assess how transient this relationship was, we assessed the relationship between hippocampal BOLD and neocortical glu/GABA ratio during the period immediately before (*a*) and after the question period (*b*). **a** Immediately prior to the question period, during presentation of the auditory cue (‘Tone’), the change in hippocampal BOLD signal between subsequently ‘remembered’ and ‘forgotten’ trials did not predict the equivalent change in glu/GABA ratio in V1. However, during the auditory cue a positive relationship was observed between the hippocampal BOLD signal and glutamate ratio in V1 (glutamate:tCr: r_16_=0.544, p=0.026; GABA:tCr: r_16_=0.28, p=0.272; glu/GABA ratio: r_16_=−0.06, p=0.809; hippocampal ROI as shown in Fig. 4a). **b** Immediately after the question period, during the inter-trial interval (‘ITI’), the change in hippocampal BOLD signal between subsequently remembered and forgotten trials did not predict the equivalent change in glu/GABA ratio in V1 (glutamate:tCr: r_16_=−0.20, p=0.444; GABA:tCr: r_16_=−0.04, p=0.869; glu/GABA ratio: r_16_=−0.04, p=0.876; hippocampal ROI as shown in Fig. 4a). **c-d** Control analyses for Fig. 4d. Upper: ROIs in parietal-occipital cortex (*c*) and brain stem (*d*), defined using a 12 mm sphere. Note: these control regions were restricted to the partial epi volume shown in Fig. 2a. Lower: Moving average showing the ratio of ‘remembered’ to ‘forgotten’ trials during the inference test: BOLD signal from parietal-occipital cortex (*c*, range [−10:10]) and brain stem (*d*, range [−4:4]) shown in grey (n=19), glutamate:tCr (red, n=19, range [− 8:8]), GABA:tCr (blue, n=19, range [−15:15]). Each point represents a 2.5s time bin (mean ± SEM).

### Supplementary Tables

**Supplementary Table 1.**
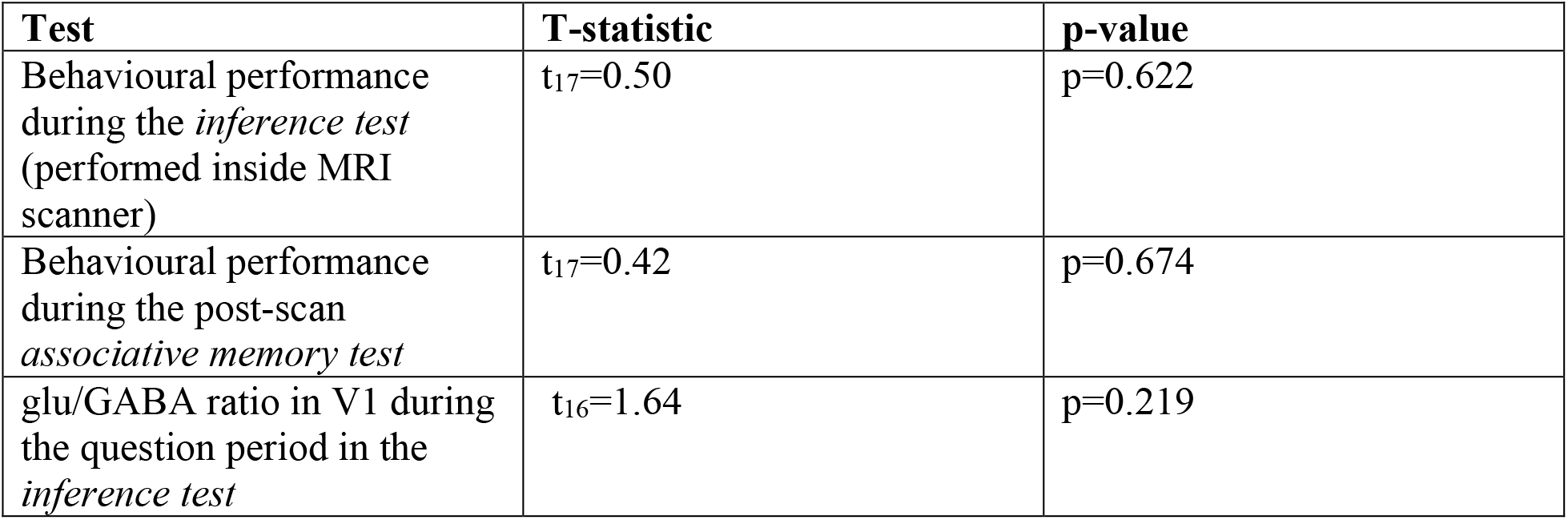
The effect of sex on behaviour and on glu/GABA ratio in V1. Using a GLM, differences in sex (male or female) were regressed onto behavioural performance during both the inference test and associative memory test, and onto glu/GABA ratio during the question period of the inference test. No significant effect of sex was observed.

| Test | T-statistic | p-value |
| --- | --- | --- |
| Behavioural performance during the *inference test* (performed inside MRI scanner) | $t_{17}=0.50$ | $p=0.622$ |
| Behavioural performance during the post-scan *associative memory test* | $t_{17}=0.42$ | $p=0.674$ |
| glu/GABA ratio in V1 during the question period in the *inference test* | $t_{16}=1.64$ | $p=0.219$ |

**Supplementary Table 2.** Number of trials per condition. The number of trials per condition, reported as mean ± SEM.

| Categorization criteria | No. of trials per condition |  |
| --- | --- | --- |
|  | 'Remembered' | 'Forgotten' |
| Using performance on both the inference test and post-scan memory test: 'correctly inferred & recalled' vs 'incorrectly inferred \| not recalled' (see Fig. 2b) | $38.95 \pm 1.56$ | $39.32 \pm 1.61$ |
| Using performance on inference test alone: 'correctly inferred' vs 'incorrectly inferred' | $59.74 \pm 1.07$ | $18.53 \pm 1.04$ |
| Using performance on post-scan memory test alone: 'recalled' vs 'not recalled' | $42.84 \pm 1.46$ | $25.53 \pm 1.51$ |

**Supplementary Table 3.** The fMRI contrast for ‘remembered’ –‘forgotten’. The fMRI BOLD signal was assessed for a contrast comparing ‘remembered’ and ‘forgotten’ trials (Fig. 2b) in the inference test. Brain regions that survived whole-volume correction for multiple comparisons are listed (p<0.05 with whole-brain FWE correction at the cluster-level).

| Brain region | P FWE-corr, cluster level | T | Peak coordinate in cluster |  |  |
| --- | --- | --- | --- | --- | --- |
|  |  |  | x | y | z |
| Left hippocampus | $P=0.017$ | 4.36 | -28 | -12 | -14 |
| Visual cortex | $P<0.001$ | 6.93 | -26 | -56 | -16 |
| Right auditory cortex | $P<0.001$ | 5.58 | 38 | -26 | 16 |
| Left posterior parietal cortex | $P<0.001$ | 5.00 | -44 | -52 | 22 |

**Supplementary Table 4.**
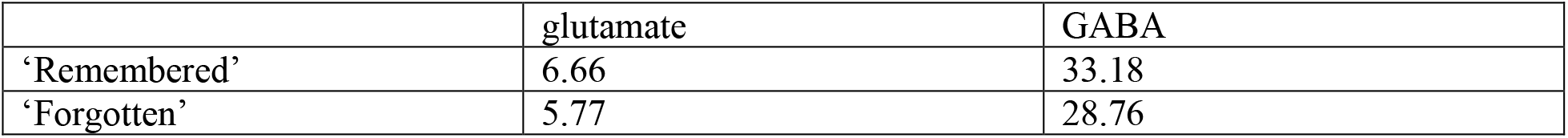
Inter-subject covariance of glutamate and GABA. Inter-subject covariances (%) for the key metabolite measurements during the ‘Question’ period of inference trials (presented in Fig. 3c–d).

|  | glutamate | GABA |
| --- | --- | --- |
| ‘Remembered’ | 6.66 | 33.18 |
| ‘Forgotten’ | 5.77 | 28.76 |

**Supplementary Table 5.** Average number of spectra (NEX) The number of spectra contributing to metabolite estimates during the various trial periods in the inference test (mean ± SEM).

|  | Tone | Question | ITI |
| --- | --- | --- | --- |
| ‘Remembered’ | 60.05±2.73 | 16.17±1.08 | 35.53±1.76 |
| ‘Forgotten’ | 47.42±2.23 | 14.61±1.21 | 28.37±1.49 |

**Supplementary Table 6.**
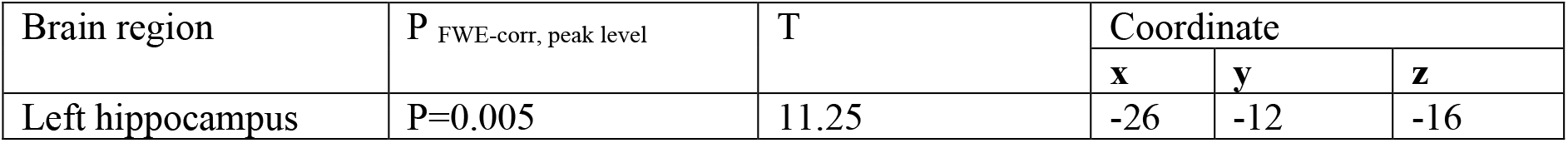
Covariance between hippocampal BOLD signal and fMRS for remembered vs. forgotten. The relationship between fMRI and fMRS during ‘remembered’ versus ‘forgotten’ trials in the inference test was assessed. To this end, fMRS measures of glu/GABA ratio from V1 for ‘remembered’ –‘forgotten’ were included as covariates in a group analysis for the equivalent fMRI contrast (p<0.05 with FWE correction at the cluster-level). The only brain region to survive whole-brain correction for multiple comparisons was the left hippocampus. Thus, the BOLD signal in left hippocampus significantly predicted individual differences in glu/GABA ratio measured from V1 during memory recall.

| Brain region | P<sub>FWE-corr, peak level</sub> | T | Coordinate |  |  |
| --- | --- | --- | --- | --- | --- |
|  |  |  | **x** | **y** | **z** |
| Left hippocampus | P=0.005 | 11.25 | -26 | -12 | -16 |

## Notes

### Competing Interest Statement

The authors have declared no competing interest.

## REFERENCES

1. Josselyn, S. A. & Tonegawa, S. Memory engrams: Recalling the past and imagining the future. Science 367, (2020).

2. Buzsáki, G. Neural syntax: cell assemblies, synapsembles and readers. Neuron 68, 362–385 (2010).

3. Squire, L. R. Memory and the hippocampus: A synthesis from findings with rats, monkeys, and humans. Psychol. Rev. 99(2), 195–231 (1992).

4. Felleman, D. J. & Essen, D. C. V. Distributed hierarchical processing in the primate cerebral cortex. Cereb. Cortex 1, 1–47 (1991).

5. Witter, M. P. Organization of the entorhinal—hippocampal system: A review of current anatomical data. Hippocampus 3, 33–44 (1993).

6. Witter, M. P., Groenewegen, H. J., Lopes da Silva, F. H. & Lohman, A. H. Functional organization of the extrinsic and intrinsic circuitry of the parahippocampal region. Prog. Neurobiol. 33, 161–253 (1989).

7. Goode, T. D., Tanaka, K. Z., Sahay, A. & McHugh, T. J. An Integrated Index: Engrams, Place Cells, and Hippocampal Memory. Neuron 107, 805–820 (2020).

8. Teyler, T. J. & DiScenna, P. The role of hippocampus in memory: A hypothesis. Neurosci. Biobehav. Rev. 9, 377–389 (1985).

9. Teyler, T. J. & Rudy, J. W. The hippocampal indexing theory and episodic memory: updating the index. Hippocampus 17, 1158–1169 (2007).

10. Pacheco Estefan, D. et al. Coordinated representational reinstatement in the human hippocampus and lateral temporal cortex during episodic memory retrieval. Nat. Commun. 10, 2255 (2019).

11. Letzkus, J. J., Wolff, S. B. E. & Lüthi, A. Disinhibition, a Circuit Mechanism for Associative Learning and Memory. Neuron 88, 264–276 (2015).

12. Barron, H. C., Auksztulewicz, R. & Friston, K. Prediction and memory: A predictive coding account. Prog. Neurobiol. 192, 101821 (2020).

13. Zhang, S. et al. Long-range and local circuits for top-down modulation of visual cortex processing. (2014).

14. Wehr, M. & Zador, A. M. Balanced inhibition underlies tuning and sharpens spike timing in auditory cortex. Nature 426(6965), 442–446 (2003).

15. Okun, M. & Lampl, I. Instantaneous correlation of excitation and inhibition during ongoing and sensory-evoked activities. Nat. Neurosci. 11, 535–537 (2008).

16. Haider, B., Duque, A., Hasenstaub, A. R. & McCormick, D. A. Neocortical network activity in vivo is generated through a dynamic balance of excitation and inhibition. J. Neurosci. 26, 4535–4545 (2006).

17. Field, R. E. et al. Heterosynaptic Plasticity Determines the Set Point for Cortical Excitatory-Inhibitory Balance. Neuron 106, 842–854 (2020).

18. McCormick, D. A., Shu, Y.-S. & Hasenstaub, A. Balanced Recurrent Excitation and Inhibition in Local Cortical Networks. in Excitatory-Inhibitory Balance: Synapses, Circuits, Systems (eds. Hensch, T. K. & Fagiolini, M.) 113–124 (Springer US, 2004). doi:10.1007/978-1-4615-0039-1_8.

19. De Graaf, R. A. In Vivo NMR Spectroscopy: Principles and Techniques. (John Wiley & Sons, 2019).

20. Mangia, S., Giove, F. & DiNuzzo, M. Metabolic Pathways and Activity-Dependent Modulation of Glutamate Concentration in the Human Brain. Neurochem. Res. 37, 2554–2561 (2012).

21. Magistretti, P. J. & Allaman, I. A cellular perspective on brain energy metabolism and functional imaging. Neuron 86, 883–901 (2015).

22. Bak, L. K., Schousboe, A. & Waagepetersen, H. S. The glutamate/GABA-glutamine cycle: aspects of transport, neurotransmitter homeostasis and ammonia transfer. J. Neurochem. 98, 641–653 (2006).

23. Rothman, D. L., Behar, K. L., Hyder, F. & Shulman, R. G. In vivo NMR Studies of the Glutamate Neurotransmitter Flux and Neuroenergetics: Implications for Brain Function. Annu. Rev. Physiol. 65, 401–427 (2003).

24. Shen, J. et al. Determination of the rate of the glutamate/glutamine cycle in the human brain by in vivo 13C NMR. Proc. Natl. Acad. Sci. 96, 8235–8240 (1999).

25. Sibson, N. R. et al. Stoichiometric coupling of brain glucose metabolism and glutamatergic neuronal activity. Proc. Natl. Acad. Sci. 95, 316–321 (1998).

26. Castro-Alamancos, M. A., Donoghue, J. P. & Connors, B. W. Different forms of synaptic plasticity in somatosensory and motor areas of the neocortex. J. Neurosci. 15, 5324–5333 (1995).

27. Trepel, C. & Racine, R. J. GABAergic modulation of neocortical long-term potentiation in the freely moving rat. Synapse 35, 120–128 (2000).

28. Floyer-Lea, A., Wylezinska, M., Kincses, T. & Matthews, P. M. Rapid modulation of GABA concentration in human sensorimotor cortex during motor learning. J. Neurophysiol. 95, 1639–1644 (2006).

29. Lunghi, C., Berchicci, M., Morrone, M. C. & Russo, F. D. Short-term monocular deprivation alters early components of visual evoked potentials. J. Physiol. 593, 4361–4372 (2015).

30. Barron, H. C. et al. Unmasking Latent Inhibitory Connections in Human Cortex to Reveal Dormant Cortical Memories. Neuron 90, 191–203 (2016).

31. Vallentin, D., Kosche, G., Lipkind, D. & Long, M. A. Inhibition protects acquired song segments during vocal learning in zebra finches. Science 351, 267–271 (2016).

32. Froemke, R. C., Merzenich, M. M. & Schreiner, C. E. A synaptic memory trace for cortical receptive field plasticity. Nature 450, 425–429 (2007).

33. Barron, H. C. et al. Neuronal Computation Underlying Inferential Reasoning in Humans and Mice. Cell https://doi.org/10.1016/j.cell.2020.08.035 (2020) doi:10.1016/j.cell.2020.08.035.

34. Bunsey, M. & Eichenbaum, H. Conservation of hippocampal memory function in rats and humans. Nature 379, 255–257 (1996).

35. DeVito, L. M., Kanter, B. R. & Eichenbaum, H. The hippocampus contributes to memory expression during transitive inference in mice. Hippocampus 20, 208–217 (2010).

36. Ip, I. B. et al. Combined fMRI-MRS acquires simultaneous glutamate and BOLD-fMRI signals in the human brain. NeuroImage 155, 113–119 (2017).

37. Horner, A. J., Bisby, J. A., Bush, D., Lin, W.-J. & Burgess, N. Evidence for holistic episodic recollection via hippocampal pattern completion. Nat. Commun. 6, 1–11 (2015).

38. Wimmer, G. E. & Shohamy, D. Preference by Association: How Memory Mechanisms in the Hippocampus Bias Decisions. Science 338, 270–273 (2012).

39. Shibata, K. et al. Overlearning hyperstabilizes a skill by rapidly making neurochemical processing inhibitory-dominant. Nat. Neurosci. 20, 470–475 (2017).

40. Gussew, A. et al. Time-resolved functional 1H MR spectroscopic detection of glutamate concentration changes in the brain during acute heat pain stimulation. NeuroImage 49, 1895–1902 (2010).

41. Bednařík, P. et al. Neurochemical responses to chromatic and achromatic stimuli in the human visual cortex. J. Cereb. Blood Flow Metab. 38, 347–359 (2018).

42. Prinsen, H., Graaf, R. A. de, Mason, G. F., Pelletier, D. & Juchem, C. Reproducibility measurement of glutathione, GABA, and glutamate: Towards in vivo neurochemical profiling of multiple sclerosis with MR spectroscopy at 7T. J. Magn. Reson. Imaging 45, 187–198 (2017).

43. Apšvalka, D., Gadie, A., Clemence, M. & Mullins, P. G. Event-related dynamics of glutamate and BOLD effects measured using functional magnetic resonance spectroscopy (fMRS) at 3 T in a repetition suppression paradigm. NeuroImage 118, 292–300 (2015).

44. Lee, S., Kruglikov, I., Huang, Z. J., Fishell, G. & Rudy, B. A disinhibitory circuit mediates motor integration in the somatosensory cortex. Nat. Neurosci. 16, 1662–1670 (2013).

45. Krabbe, S. et al. Adaptive disinhibitory gating by VIP interneurons permits associative learning. Nat. Neurosci. 22, 1834–1843 (2019).

46. Koolschijn, R. S. et al. The Hippocampus and Neocortical Inhibitory Engrams Protect against Memory Interference. Neuron 101, 528–541 (2019).

47. Nikolova, S., Stark, S. M. & Stark, C. E. L. 3T hippocampal glutamate-glutamine complex reflects verbal memory decline in aging. Neurobiol. Aging 54, 103–111 (2017).

48. Schmitz, T. W., Correia, M. M., Ferreira, C. S., Prescot, A. P. & Anderson, M. C. Hippocampal GABA enables inhibitory control over unwanted thoughts. Nat. Commun. 8, (2017).

49. Vaz, A. P., Wittig, J. H., Inati, S. K. & Zaghloul, K. A. Replay of cortical spiking sequences during human memory retrieval. Science 367, 1131–1134 (2020).

50. Isaacson, J. S. & Scanziani, M. How Inhibition Shapes Cortical Activity. Neuron 72, 231–243 (2011).

51. Vogels, T. P., Sprekeler, H., Zenke, F., Clopath, C. & Gerstner, W. Inhibitory Plasticity Balances Excitation and Inhibition in Sensory Pathways and Memory Networks. Science 334, 1569–1573 (2011).

52. Kuchibhotla, K. V. et al. Parallel processing by cortical inhibition enables context-dependent behavior. Nat. Neurosci. 20, 62–71 (2017).

53. Wolff, S. B. E. et al. Amygdala interneuron subtypes control fear learning through disinhibition. Nature 509, 453–458 (2014).

54. Courtin, J. et al. Prefrontal parvalbumin interneurons shape neuronal activity to drive fear expression. Nature 505, 92–96 (2014).

55. Mekle, R. et al. Detection of metabolite changes in response to a varying visual stimulation paradigm using short-TE 1H MRS at 7 T. NMR Biomed. 30, e3672 (2017).

56. Hong, D., Rankouhi, S. R., Thielen, J.-W., Asten, J. J. A. van & Norris, D. G. A comparison of sLASER and MEGA-sLASER using simultaneous interleaved acquisition for measuring GABA in the human brain at 7T. PLOS ONE 14, e0223702 (2019).

57. Lally, N. et al. Glutamatergic correlates of gamma-band oscillatory activity during cognition: A concurrent ER-MRS and EEG study. NeuroImage 85, 823–833 (2014).

58. Rae, C. A Guide to the Metabolic Pathways and Function of Metabolites Observed in Human Brain 1H Magnetic Resonance Spectra. Neurochem. Res. 39, 1–36 (2014).

59. Myers, J. F., Nutt, D. J. & Lingford-Hughes, A. R. γ-aminobutyric acid as a metabolite: Interpreting magnetic resonance spectroscopy experiments. J. Psychopharmacol. (Oxf.) 30, 422–427 (2016).

60. De Graaf, A. A. & Bovée, W. M. M. J. Improved quantification of in vivo1H NMR spectra by optimization of signal acquisition and processing and by incorporation of prior knowledge into the spectral fitting. Magn. Reson. Med. 15, 305–319 (1990).

61. Kauppinen, R. A. & Williams, S. R. Nondestructive Detection of Glutamate by 1H Nuclear Magnetic Resonance Spectroscopy in Cortical Brain Slices from the Guinea Pig: Evidence for Changes in Detectability During Severe Anoxic Insults. J. Neurochem. 57, 1136–1144 (1991).

62. Fox, P. T., Raichle, M. E., Mintun, M. A. & Dence, C. Nonoxidative glucose consumption during focal physiologic neural activity. Science 241, 462–464 (1988).

63. Fox, P. T. & Raichle, M. E. Focal physiological uncoupling of cerebral blood flow and oxidative metabolism during somatosensory stimulation in human subjects. Proc. Natl. Acad. Sci. 83, 1140–1144 (1986).

64. Gjedde, A., Marrett, S. & Vafaee, M. Oxidative and Nonoxidative Metabolism of Excited Neurons and Astrocytes. J. Cereb. Blood Flow Metab. 22, 1–14 (2002).

65. Stanley, J. A. & Raz, N. Functional Magnetic Resonance Spectroscopy: The “New” MRS for Cognitive Neuroscience and Psychiatry Research. Front. Psychiatry 9, (2018).

66. Cleve, M., Gussew, A. & Reichenbach, J. R. In vivo detection of acute pain-induced changes of GABA+ and Glx in the human brain by using functional 1H MEGA-PRESS MR spectroscopy. NeuroImage 105, 67–75 (2015).

67. Stagg, C. J., Bachtiar, V. & Johansen-Berg, H. The role of GABA in human motor learning. Curr. Biol. 21, 480–484 (2011).

68. Puts, N. A. J., Edden, R. A. E., Evans, C. J., McGlone, F. & McGonigle, D. J. Regionally Specific Human GABA Concentration Correlates with Tactile Discrimination Thresholds. J. Neurosci. 31, 16556–16560 (2011).

69. Scholl, J. et al. Excitation and inhibition in anterior cingulate predict use of past experiences. eLife 6, (2017).

70. Stagg, C. & Rothman, D. L. Magnetic Resonance Spectroscopy: Tools for Neuroscience Research and Emerging Clinical Applications. (Academic Press, 2013).

71. Jelen, L. A., King, S., Mullins, P. G. & Stone, J. M. Beyond static measures: A review of functional magnetic resonance spectroscopy and its potential to investigate dynamic glutamatergic abnormalities in schizophrenia. J. Psychopharmacol. (Oxf.) 32, 497–508 (2018).

72. Barron, H. C., Garvert, M. M. & Behrens, T. E. J. Repetition suppression: a means to index neural representations using BOLD? Philos. Trans. R. Soc. Lond. B. Biol. Sci. 371, (2016).

73. Andreychenko, A., Boer, V. O., Castro, C. S. A. de, Luijten, P. R. & Klomp, D. W. J. Efficient spectral editing at 7 T: GABA detection with MEGA-sLASER. Magn. Reson. Med. 68, 1018–1025 (2012).

74. Govindaraju, V., Young, K. & Maudsley, A. A. Proton NMR chemical shifts and coupling constants for brain metabolites. NMR Biomed. 13, 129–153 (2000).

75. Puts, N. A. J. & Edden, R. A. E. In vivo magnetic resonance spectroscopy of GABA: A methodological review. Prog. Nucl. Magn. Reson. Spectrosc. 60, 29–41 (2012).

76. Bottomley, P. A. Spatial Localization in NMR Spectroscopy in Vivo. Ann. N. Y. Acad. Sci. 508, 333–348 (1987).

77. Mescher, M., Merkle, H., Kirsch, J., Garwood, M. & Gruetter, R. Simultaneous in vivo spectral editing and water suppression. NMR Biomed. 11, 266–272 (1998).

78. Trabesinger, A. H. & Boesiger, P. Improved selectivity of double quantum coherence filtering for the detection of glutathione in the human brain in vivo. Magn. Reson. Med. 45, 708–710 (2001).

79. Terpstra, M., Marjanska, M., Henry, P.-G., Tkáč, I. & Gruetter, R. Detection of an antioxidant profile in the human brain in vivo via double editing with MEGA-PRESS. Magn. Reson. Med. 56, 1192–1199 (2006).

80. Lin, Y., Stephenson, M. C., Xin, L., Napolitano, A. & Morris, P. G. Investigating the Metabolic Changes due to Visual Stimulation using Functional Proton Magnetic Resonance Spectroscopy at 7 T. J. Cereb. Blood Flow Metab. 32, 1484–1495 (2012).

81. Robertson, C. E., Ratai, E.-M. & Kanwisher, N. Reduced GABAergic Action in the Autistic Brain. Curr. Biol. 26, 80–85 (2016).

82. Taylor, R. et al. Functional magnetic resonance spectroscopy of glutamate in schizophrenia and major depressive disorder: anterior cingulate activity during a color-word Stroop task. Npj Schizophr. 1, 1–8 (2015).

83. Preston, A. R. & Eichenbaum, H. Interplay of hippocampus and prefrontal cortex in memory. Curr. Biol. 23, R764–R773 (2013).

84. Brogden, W. J. Sensory pre-conditioning. J. Exp. Psychol. 25, 323–332 (1939).

85. Brink, W. M. & Webb, A. G. High Permittivity Pads Reduce Specific Absorption Rate, Improve B1 Homogeneity, and Increase Contrast-to-Noise Ratio for Functional Cardiac MRI at 3 T. Magn. Reson. Med. 71, 1632–1640 (2014).

86. Hess, A. T., Tisdall, M. D., Andronesi, O. C., Meintjes, E. M. & Van Der Kouwe, A. J. W. Real-Time Motion and B0 Corrected Single Voxel Spectroscopy Using Volumetric Navigators. Magn. Reson. Med. 66, 314–323 (2011).

87. Öz, G. & Tkáč, I. Short-Echo, Single-Shot, Full-Intensity Proton Magnetic Resonance Spectroscopy for Neurochemical Profiling at 4 T : Validation in the Cerebellum and Brainstem. Magn. Reson. Med. 65, 901–910 (2011).

88. Terpstra, M. et al. Test-retest reproducibility of neurochemical profiles with short-echo, single-voxel MR spectroscopy at 3T and 7T. Magn. Reson. Med. 76, 1083–1091 (2016).

89. Provencher, S. W. Estimation of Metabolite Concentrations from Localized in Vivo Proton NMR Spectra. Magn. Reson. Med. 30, 672–679 (1993).

90. Worsley, K. J. & Friston, K. J. Analysis of fMRI time-series revisited--again. NeuroImage 2, 173–181 (1995).

91. Mangia, S. et al. Sensitivity of single-voxel 1H-MRS in investigating the metabolism of the activated human visual cortex at 7 T. Magn. Reson. Imaging 24, 343–348 (2006).

